# High basal autophagic activity in the brain revealed by systemic quantitative analysis using GFP-LC3-RFP mice

**DOI:** 10.64898/2026.05.20.726446

**Authors:** Yuki Kanda, Tomoya Eguchi, Hideaki Morishita, Yutaro Hama, Manabu Abe, Kenji Sakimura, Noboru Mizushima

## Abstract

Autophagy is a fundamental intracellular degradation pathway with vital physiological functions. Although it is well known that autophagy is activated during starvation, the extent of basal autophagy remains unclear owing to challenges in measuring autophagic flux *in vivo*. In this study, we developed autophagy reporter (GFP–LC3–RFP) mice and quantified basal autophagic flux across tissues by comparing normal and autophagy-deficient conditions. Comparative analyses revealed uniformly low basal autophagic flux during embryogenesis, but significant tissue-specific variation in adult mice. In contrast to previous assumptions that basal autophagy in the brain is low, the brain, along with the liver and kidney, exhibited higher basal autophagic flux than the heart, skeletal muscle, and intestine. These data serve as foundational information on basal autophagic flux in mammals and provide a plausible explanation for the severe neurological phenotypes linked to autophagy gene mutations in mice and humans.

## Introduction

Macroautophagy (hereafter referred to as autophagy), the process by which cytosolic materials are transported to lysosomes, is one of the primary intracellular degradation mechanisms widely conserved among eukaryotes (Dikic & Elazar, 2018; Yamamoto et al., 2023). Autophagy occurs constitutively at low levels under basal conditions and is activated in response to stimuli that include nutrient depletion, organelle damage, and pathogen invasion. It plays crucial roles in maintaining cellular homeostasis by supplying degradation products and eliminating unwanted intracellular components (Mizushima & Komatsu, 2011). Recent studies have shown that impaired autophagy is linked to various diseases, including neurodevelopmental and neurodegenerative disorders (Klionsky, Petroni, et al., 2021; Mizushima & Levine, 2020), and has been implicated in aging (Hansen et al., 2018; Leidal et al., 2018).

Brain autophagy levels are generally considered low, given that autophagosomes are rarely observed under basal conditions (Kuma et al., 2004; Nixon et al., 2005) and the brain shows a limited autophagic response to starvation(Kuma et al., 2004). However, static snapshots of autophagosome counts do not necessarily reflect the actual flux or degradative activity(Klionsky, Abdel-Aziz, et al., 2021). Several methods to monitor autophagic flux *in vivo* have been developed, including LC3 turnover assays using lysosomal inhibitors (Haspel et al., 2011) and tandem fluorescent LC3 (tfLC3) reporters (Castillo et al., 2013; Hariharan et al., 2011; Lee et al., 2019; Li et al., 2014; Oliva Trejo et al., 2020). Although conceptually straightforward, inhibitor-based methods have not been applied to the brain, likely as a consequence of the poor delivery of inhibitors to the nervous system. Although tfLC3 reporters have been used to visualize basal autophagic activity in the brain with intra-cerebroventricular infusion of inhibitors (Lee et al., 2019), cross-organ comparisons have not been performed. Consequently, the actual extent of brain autophagic flux has remained elusive.

Mouse and human genetic studies have suggested that basal autophagy in the brain is particularly important, implying that its true activity has been substantially underestimated. Brain-specific deletion of autophagy-related genes, including *Atg5*, *Atg7* and *Fip200*, leads to severe phenotypes, including neurodegeneration, motor dysfunction, and premature death within a few months (Hara et al., 2006; Komatsu et al., 2006; Komatsu et al., 2007; Liang et al., 2010). Furthermore, mutations in core autophagy-related genes, including *ATG5*, *ATG7*, *WIPI2*, *WDR45B/WIPI3*, and *WDR45*/*WIPI4*, have been identified as the causes of human neurodevelopmental and neurodegenerative diseases (Almannai et al., 2022; Collier et al., 2021; Haack et al., 2012; Jelani et al., 2019; Kim et al., 2016; Maroofian et al., 2021; Saitsu et al., 2013; Suleiman et al., 2018). These findings suggest that basal autophagic flux in the brain might be higher than previously thought, although the actual level remains uncertain owing to the lack of quantitative cross-tissue comparisons.

To quantitatively assess basal autophagic flux across different tissues, including the brain, we newly generated a knock-in mouse line that ubiquitously expresses the GFP–LC3–RFP autophagic flux reporter (Kaizuka et al., 2016). Using this mouse line, we conducted a comparative analysis of starvation-induced and basal autophagic flux across major tissues in embryos and adult mice. We determined basal autophagy as the difference in the GFP/RFP signal ratio between normal and autophagy-deficient conditions. We found that basal autophagic flux varies substantially among adult tissues, whereas it was uniformly low during embryogenesis. Notably, adult mice exhibited high basal autophagic flux in the brain, as well as in the liver and kidney. Our findings provide direct quantitative evidence that contradicts the long-standing assumption that basal autophagy is low in the brain.

## Results

### Generation of a knock-in mouse line systemically expressing the GFP–LC3–RFP reporter

We previously developed a transgenic mouse line expressing GFP–LC3–RFP–LC3ΔG, but reporter expression was restricted mainly to the skeletal muscle (Kaizuka et al., 2016). To resolve this key limitation, we aimed to generate a knock-in mouse line with ubiquitous expression of the autophagic flux reporter by inserting the reporter gene into the *Rosa26* (reverse orientation splice acceptor 26) locus, which enables ubiquitous transgene expression across developmental stages without disrupting endogenous gene function (Soriano, 1999; Zambrowicz et al., 1997) (Figure 1A). A previous study demonstrated that repeated insertion of a transgene at the same locus can exert a repressive effect on its expression (Garrick et al., 1998). To avoid this issue, GFP–LC3–RFP (without LC3ΔG) was used as autophagic flux reporter, which we previously validated as functionally equivalent to GFP–LC3–RFP–LC3ΔG (Kaizuka et al., 2016). Upon expression in cells, the reporter is cleaved by the endogenous protease ATG4, generating equimolar amounts of GFP–LC3 and RFP. GFP–LC3 undergoes autophagic degradation, while RFP remains in the cytosol. Therefore, the GFP/RFP signal ratio is inversely correlated with autophagic flux (Figure 1B).

**Figure 1.**
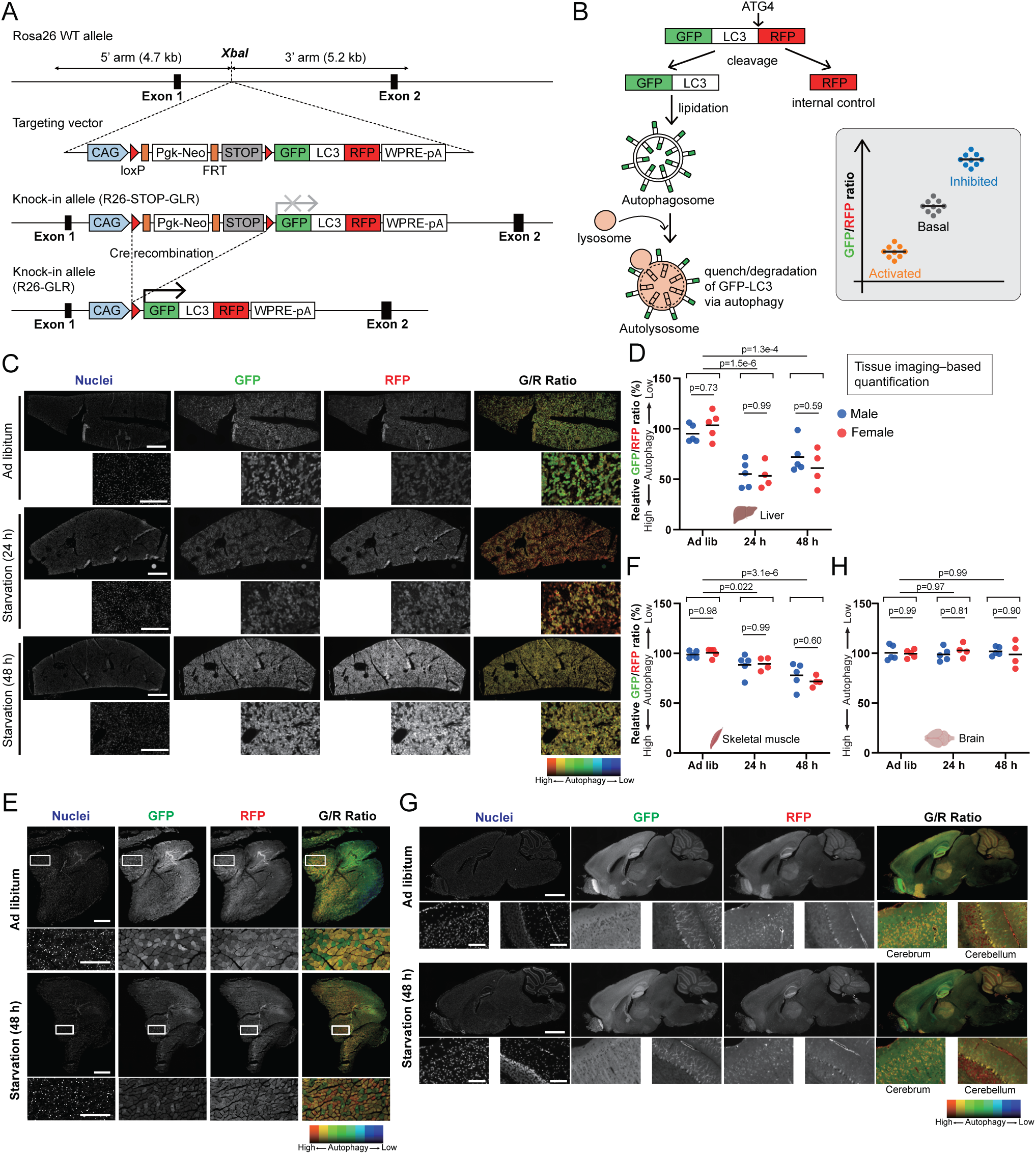
Generation of GFP–LC3–RFP knock-in mice and the detection of starvation-induced autophagy. (A) Schematic illustration of the knock-in strategy to generate R26-STOP-GLR mice expressing the GFP–LC3–RFP autophagic flux reporter. The cassette containing the CAG promoter, a loxP-flanked STOP cassette, and the GFP–LC3–RFP reporter was inserted into the intron between exons 1 and 2 of the *Rosa26* locus. (B) Schematic illustration of the principle of the GFP–LC3–RFP reporter to evaluate autophagic flux. (C, E, G) Representative fluorescence and corresponding GFP/RFP ratio images of the liver (C), skeletal muscle (E), and brain (G). Mice (3–4-month-old) were starved for 24 h or 48 h. Scale bars, 1 mm (low magnification) and 200 µm (high magnification). (D, F, H) Quantification of the GFP/RFP ratio in the liver (D), skeletal muscle (F), and brain (H). Each dot represents an individual mouse; bars indicate mean values; *n* = 4–5 per sex per condition, total *n* = 9–10 per group. Data were analyzed by two-way ANOVA with Sidak’s and Tukey’s post hoc tests for sex and starvation comparisons, respectively.

We generated Rosa26-CAG-loxP-STOP-loxP-GFP-LC3-RFP knock-in mice (hereafter referred to as R26-STOP-GLR), in which expression of the reporter is controlled by the cytomegalovirus early enhancer/chicken β-actin (CAG) promoter and a lox-stop-lox (LSL) cassette, enabling Cre-dependent expression. To confirm conditional reporter expression, R26-STOP-GLR mice were crossed with Vasa-Cre transgenic mice, which specifically express Cre recombinase in germ cells (Gallardo et al., 2007). Successful removal of the STOP cassette—resulting in the generation of R26-GLR mice (Rosa26-CAG-GFP-LC3-RFP)—was confirmed by genomic PCR (Figure1—figure supplement 1A). Both R26-STOP-GLR and R26-GLR mice developed normally and were fertile. GFP–LC3 and RFP were systemically detected, whereas the full length of GFP–LC3–RFP was not detected, indicating that GFP–LC3–RFP was ubiquitously expressed and completely cleaved by endogenous ATG4 proteases (Figure S1B). Additionally, reporter expression did not alter p62 levels or LC3 lipidation, suggesting that it did not perturb endogenous autophagic flux (Figure1—figure supplement 1B and Figure 1—figure supplement 3). Imaging of tissue sections showed clear GFP and RFP signals in R26-GLR mice, but not in wild-type (WT) or R26-STOP-GLR mice (Figure1—figure supplement 1C and Figure 1—figure supplement 2). The liver showed mosaic reporter expression, although all other tissues analyzed exhibited ubiquitous expression (Figure1—figure supplement 1C and Figure 1—figure supplement 2). This is consistent with a previous report showing that CAG promoter-driven transgene expression from the Rosa26 locus resulted in mosaic expression in hepatocytes (Tchorz et al., 2012). These results demonstrate that R26-GLR mice systemically express the autophagic flux reporter GFP–LC3–RFP without interfering with endogenous autophagy.

### Detection of starvation-induced autophagy by tissue imaging

To determine whether the R26-GLR mouse line can be used to monitor autophagic flux *in vivo*, we first analyzed changes in the GFP/RFP ratio in response to starvation. R26-GLR mice (3–4-month-old) were starved for 24 or 48 h. The GFP/RFP ratio was calculated from imaging of tissue sections by averaging the pixel-wise GFP/RFP ratios within the tissue area. To highlight regions with altered autophagic flux, the ratio images were visualized using a blue-to-red pseudo-color scale, in which blue indicates lower autophagic flux (higher GFP/RFP ratio) and red indicates higher autophagic flux (lower GFP/RFP ratio). Tissue imaging showed a decrease in the GFP/RFP ratio in the liver and skeletal muscle after 24 h of starvation, with a continued decline in the muscle at 48 h, showing comparable response between male and female mice (Figure 1C–F). These results are consistent with previously reported findings using GFP–LC3–RFP–LC3ΔG transgenic mice (Kaizuka et al., 2016) and pHluorin–LC3–mCherry KI mice (Aoyama et al., 2023). Immunoblotting also showed a decreased GFP/RFP ratio in the starved liver and skeletal muscle (Figure1—figure supplement 1D, E, G, H), indicating that the reduction in GFP signal was caused primarily by increased autophagic degradation of GFP–LC3 rather than quenching of GFP in acidic autolysosomes.

In the liver, mTORC1 activity, as assessed by the phosphorylation status of ribosomal protein S6 kinase beta-1 (S6K1) and eukaryotic translation initiation factor 4E-binding protein 1 (4E-BP1), was suppressed after a 24-h starvation period and had partially recovered by 48 h (Figure1—figure supplement 1D). In the skeletal muscle, mTORC1 inhibition persisted at both time points (Figure1—figure supplement 1E). LC3-II levels also increased, reflecting up-regulation of autophagy upon starvation. Taken together, these results suggest that R26-GLR mice can detect starvation-induced autophagy in both liver and skeletal muscle. In contrast, tissue imaging revealed an unchanged GFP/RFP ratio in the brain after 24 and 48 h of starvation in both sexes (Figure 1G, H). Immunoblotting also showed a stable GFP/RFP ratio (Figure1—figure supplement 1F, I). Furthermore, mTORC1 activity and LC3-II levels were unaffected in the brain (Figure1—figure supplement 1F), indicating that brain autophagy is resistant to starvation, consistent with previous reports (Kaade et al., 2024; Mizushima et al., 2004). Collectively, these findings demonstrate that R26-GLR mice provide a versatile *in vivo* model for evaluating autophagic flux.

### Detection of starvation-induced autophagy by a microplate reader

Although the calculation of the GFP/RFP ratio through tissue section imaging provides valuable spatial information about autophagic flux, this method is time-consuming and labor-intensive. Therefore, we developed a simplified approach for measuring the GFP/RFP ratio using a microplate reader (Figure 2A). GFP and RFP signals were robustly detected in lysates from the brain, liver, and kidney of R26-GLR mice, with good linearity of the signals (Figure 2B, Figure 2—figure supplement 1A). Both GFP and RFP signals were detectable even using only 200 µL of 6 mg tissue/mL lysate (corresponding to 1.2 mg of tissue). At high tissue lysate concentrations (e.g., 100 mg/mL), the signals slightly deviated from linearity, possibly owing to sample opacity interfering with signal detection. Therefore, GFP and RFP signals were measured at a concentration of 25 mg tissue/mL in all organs, demonstrating that both signals were easily detected, with high signal-to-noise (S/N) ratios (Figure 2C). We confirmed that signal detection was not substantially affected by variation in lysis buffer composition or centrifugation speed (Figure 2—figure supplement 1B–E) and that the majority of the signal resided in the soluble fraction (Figure 2—figure supplement 1F, G). Thus, we adopted a standardized protocol using Triton X-100 and low-speed centrifugation for subsequent measurements.

**Figure 2.**
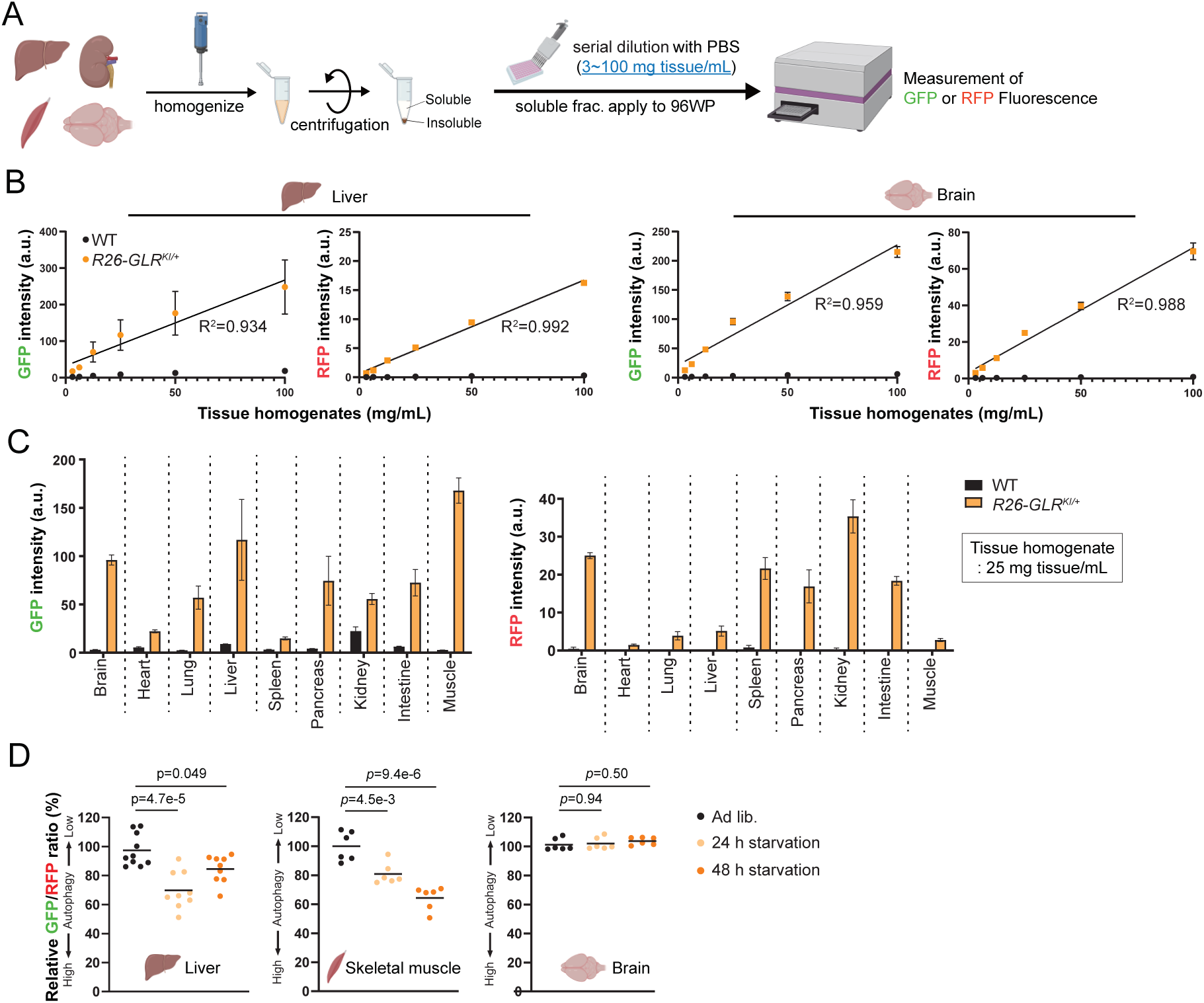
A simple, rapid, and semi-high-throughput measurement of the GFP/RFP ratio using a microplate reader. (A) Schematic illustration of the microplate reader-based assay detecting GFP–LC3–RFP reporter signals. Created with BioRender.com. (B) Soluble fractions from liver and brain lysates of WT and R26-GLR mice (3–4-month-old) were serially diluted and measured using a microplate reader (*n* = 3 per group). Data are presented as mean ± SD values. The coefficients of determination (*R*²) from linear regressions of measurements of tissue concentration and GFP signal for R26-GLR mice are shown. (C) Soluble fractions from tissue lysates of WT and R26-GLR mice (3–4-month-old) were diluted 4× and measured using a microplate reader (*n* = 3 per group). Bars represent mean ± SD values. (D) GFP/RFP ratios in the liver, skeletal muscle, and brain under 24- or 48-h starvation conditions, calculated from microplate reader measurements. Each dot represents an individual mouse; bars indicate mean values; *n* = 6–9 per group. Differences among groups were analyzed by one-way ANOVA with Tukey’s post hoc test.

We then evaluated whether autophagic flux could be assessed via this microplate reader-based method at a level comparable to that obtained by tissue section imaging. We prepared 24- and 48-h-starved mice, and their tissue lysates were subjected to the microplate reader assay. Upon starvation, the GFP/RFP ratio decreased markedly in both liver and skeletal muscle at 24 and 48 h (Figure 2D). In contrast, the ratio remained unchanged in the brain at either time point (Figure 2E). These results were consistent with those obtained by tissue section imaging (Figure 1D, F, H). Thus, this simple plate reader-based method represents a substantial improvement in throughput for measuring autophagic flux *in vivo*. To evaluate autophagic flux comprehensively, we employed both tissue imaging and the plate reader-based method in subsequent experiments.

### Comparative assessment of basal autophagic flux among and within tissues in *embryos*

Next, we investigated whether basal autophagic flux varies across and within embryonic tissues. We assumed that the difference in GFP/RFP ratios between WT and autophagy-deficient embryos reflected the basal autophagic activity in each tissue. To generate autophagy-deficient embryos, Atg5—an essential gene for autophagy and lipidation of ATG8 family proteins (Mizushima et al., 1998; Mizushima et al., 2001)—was conditionally deleted. Atg5 flox mice (Hara et al., 2006) were crossed with Vasa-Cre mice (Gallardo et al., 2007) to obtain mice with the Atg5-deleted allele (*Atg5^Δ^*), which were then mated with R26-GLR (*R26-GLR^KI^*) mice to generate embryos with the genotypes *R26-GLR^KI/+^;Atg5^+/+^*, *R26-GLR^KI/+^;Atg5^Δ/+^*, and *R26-GLR^KI/+^;Atg5^Δ/Δ^*, hereafter referred to as WT, Atg5 HT (heterozygous), and Atg5 KO, respectively (Figure 3A). Embryos were harvested at embryonic day 19.5 (E19.5) and decapitated to minimize blood-derived interference (Figure 3A). Loss of ATG5 and LC3-II, along with accumulation of p62, were observed in Atg5 KO embryos, while Atg5 HT embryos showed levels comparable to those of WT embryos (Figure 3—figure supplement 1A). GFP–LC3 levels increased only in Atg5 KO embryos, while RFP levels remained unchanged (Figure 3—figure supplement 1A). GFP and RFP signals of embryonic tissue lysates were robustly detected by a microplate reader, revealing S/N ratios comparable to those of adult tissues (Figure 3—figure supplement 1B). Consistent with immunoblotting, the GFP/RFP ratios were similar between WT and Atg5 HT embryos, whereas Atg5 KO embryos exhibited an approximately 1.5-fold increase in both the head and body (Figure 3B).

**Figure 3.**
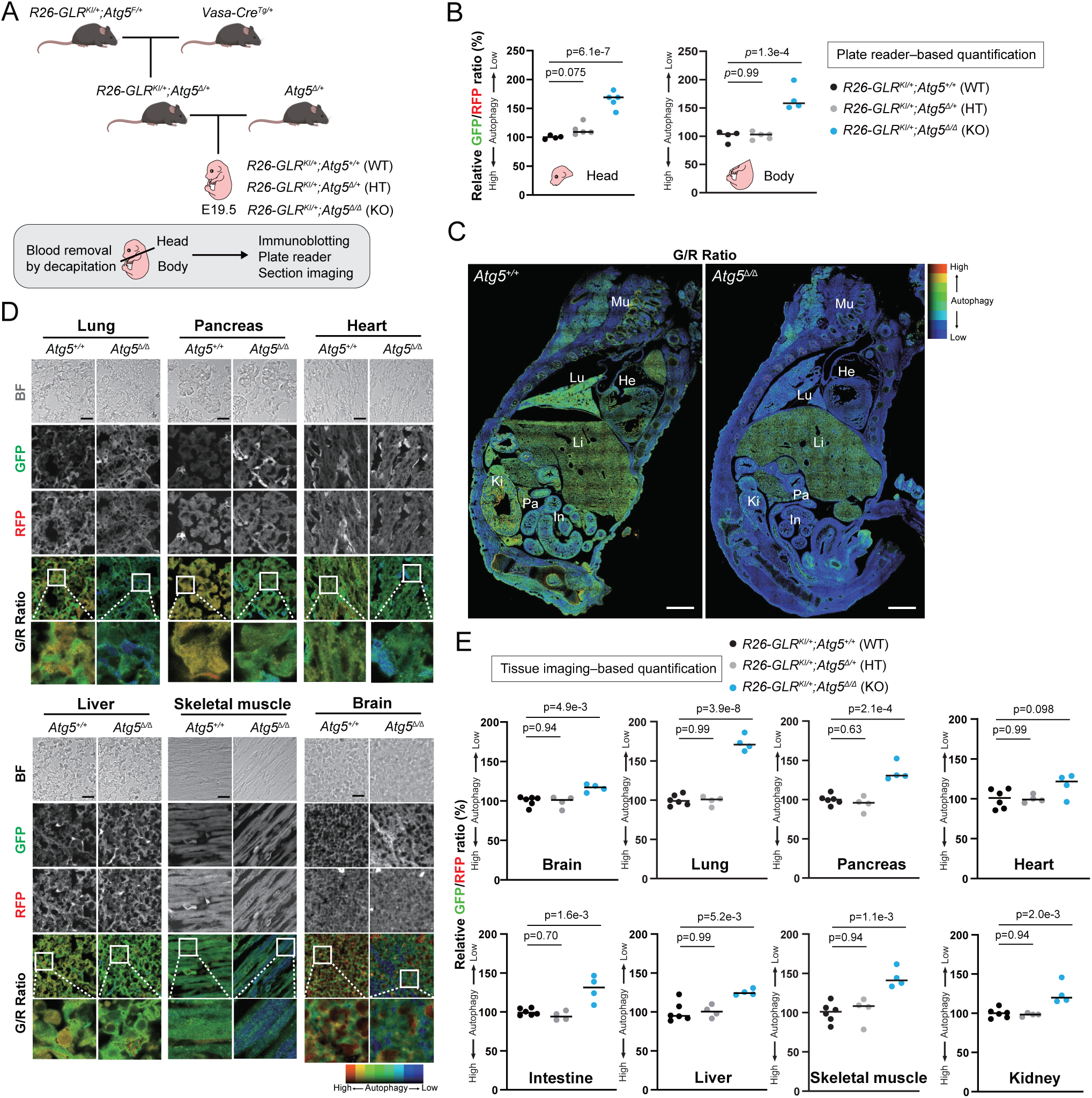
Uniform basal autophagic flux across multiple tissues during embryonic development. (A) Schematic illustration of the breeding strategy for obtaining GFP–LC3–RFP–expressing Atg5 knockout embryos (E19.5). Embryo dissection and detection were performed as described. Created with BioRender.com. (B) GFP/RFP ratios in Atg5 wild-type (WT), heterozygous (HT), or knockout (KO) embryos measured using a microplate reader. Each dot represents an individual embryo; bars indicate mean values; *n* = 4–5 per group. Differences among groups were analyzed by one-way ANOVA with Tukey’s post hoc test. (C) Representative GFP/RFP ratio images of the body of *Atg5* WT and *Atg5* KO embryos. Scale bar, 1 mm; Mu, skeletal muscle; He, heart; Lu, lung; Li, liver; Pa, pancreas; Ki, kidney; and In, intestine. (D) Representative fluorescence and corresponding GFP/RFP ratio images of major tissues of *Atg5* WT and *Atg5* KO embryos. Scale bar, 100 µm. (E) The GFP/RFP ratio calculated from tissue imaging of *Atg5* WT, Atg5 HT, and *Atg5* KO embryos. Each dot represents an individual embryo; bars indicate mean values; *n* = 4–6 per group. Differences among groups were analyzed by one-way ANOVA with Tukey’s post hoc test.

To obtain spatial resolution information, tissue imaging was also performed. Despite lower S/N ratios compared to adult tissues, reporter expression was clearly detectable in R26-GLR embryos (Figure 3—figure supplement 1C). Whole-section imaging revealed an elevated GFP/RFP ratio throughout the Atg5 KO embryos, as visualized by a pseudo-color shift toward blue (Figure 3C, Figure 3—figure supplement 1D). High-magnification analysis of individual organs revealed a 1.2- to 1.5-fold increase in GFP/RFP ratios, consistent with the plate reader-based assay (Figure 3D, E). These findings suggest that basal autophagic flux is generally comparable across embryonic tissues, with only subtle differences observed among individual tissues.

In addition to tissue-level analyses, we also assessed basal autophagic flux at the single-cell level. At this developmental stage, glomeruli exhibit mature morphology comparable to those in adult mice and are composed primarily of podocytes, mesangial cells, and epithelial cells (Schell et al., 2014). In WT embryos, the GFP/RFP ratio was lower in podocytes than in the surrounding glomerular cells. However, this cell-type-specific difference was abolished in Atg5 KO embryos (Figure 3—figure supplement 1E). This result suggests that podocytes displayed higher basal autophagic flux compared with neighboring glomerular cells and demonstrates that R26-GLR mice can be used to assay differences in basal autophagic flux not only among tissues but also among individual cell types.

### Basal autophagic flux across tissues and neuronal cell populations in adult mice

We next examined basal autophagic flux across tissues in adult mice. To this end, we utilized tamoxifen-inducible Atg5-deficient mice harboring the GFP–LC3–RFP reporter, because systemic deletion of *Atg5* is lethal. These mice carried the floxed *Atg5* allele (*Atg5^F^*) and a *CreERT2* transgene (*CreERT2^Tg^*) under the control of the human ubiquitin C (UBC) promoter (Ruzankina et al., 2007). *R26-GLR^KI/+^;Atg5^+/+^*, *R26-GLR^KI/+^;Atg5^flox/flox^*, *R26-GLR^KI/+^;Atg5^flox/+^;CreERT2^Tg/+^*, and *R26-GLR^KI/+^;Atg5^flox/flox^;CreERT2^Tg/+^* mice are hereafter referred to as WT, *Atg5^F/F^*, *Atg5^F/+^;Cre*, and *Atg5^F/F^;Cre*, respectively. Tamoxifen was administered once weekly for four weeks, and tissues were collected four weeks after the final injection (Figure 4A). Cre-mediated recombination was confirmed by genomic PCR (Figure 4—figure supplement 1A, B). *Atg5^F/F^* mice that did not receive tamoxifen injection already showed reduced ATG5 expression but did not affect LC3 lipidation and p62 levels in the brain and kidney (Figure 4B, Figure 4—figure supplement 1C), indicating that the floxed Atg5 allele is hypomorphic (as expected given that the neo cassette was inserted into the third intron and retained (Hara et al., 2006)), consistent with a previous report (Lin et al., 2014), yet retains functional autophagy. Similarly, *Atg5^F/+^;Cre* mice treated with tamoxifen exhibited a modest reduction in ATG5 expression, with no significant changes in LC3 lipidation or p62 levels (Figure 4B, Figure 4—figure supplement 1C), supporting the inference that heterozygous *Atg5* loss does not compromise autophagy under basal conditions. In contrast, tamoxifen-treated *Atg5^F/F^;Cre* mice displayed a profound reduction in ATG5 expression in both the brain and kidney, accompanied by the accumulation of LC3-I and p62 and a marked reduction in LC3-II, indicative of impaired autophagic flux (Figure 4B). This pattern was recapitulated in most other tissues, even in the pancreas, which showed substantial but incomplete ATG5 deletion (Figure 4—figure supplement 1D, E). These results demonstrate that tamoxifen administration efficiently induced *Atg5* deletion across a wide range of adult tissues (Figure 4—figure supplement 1D, E), enabling the assessment of basal autophagic flux under near-complete autophagy-deficient conditions.

**Figure 4.**
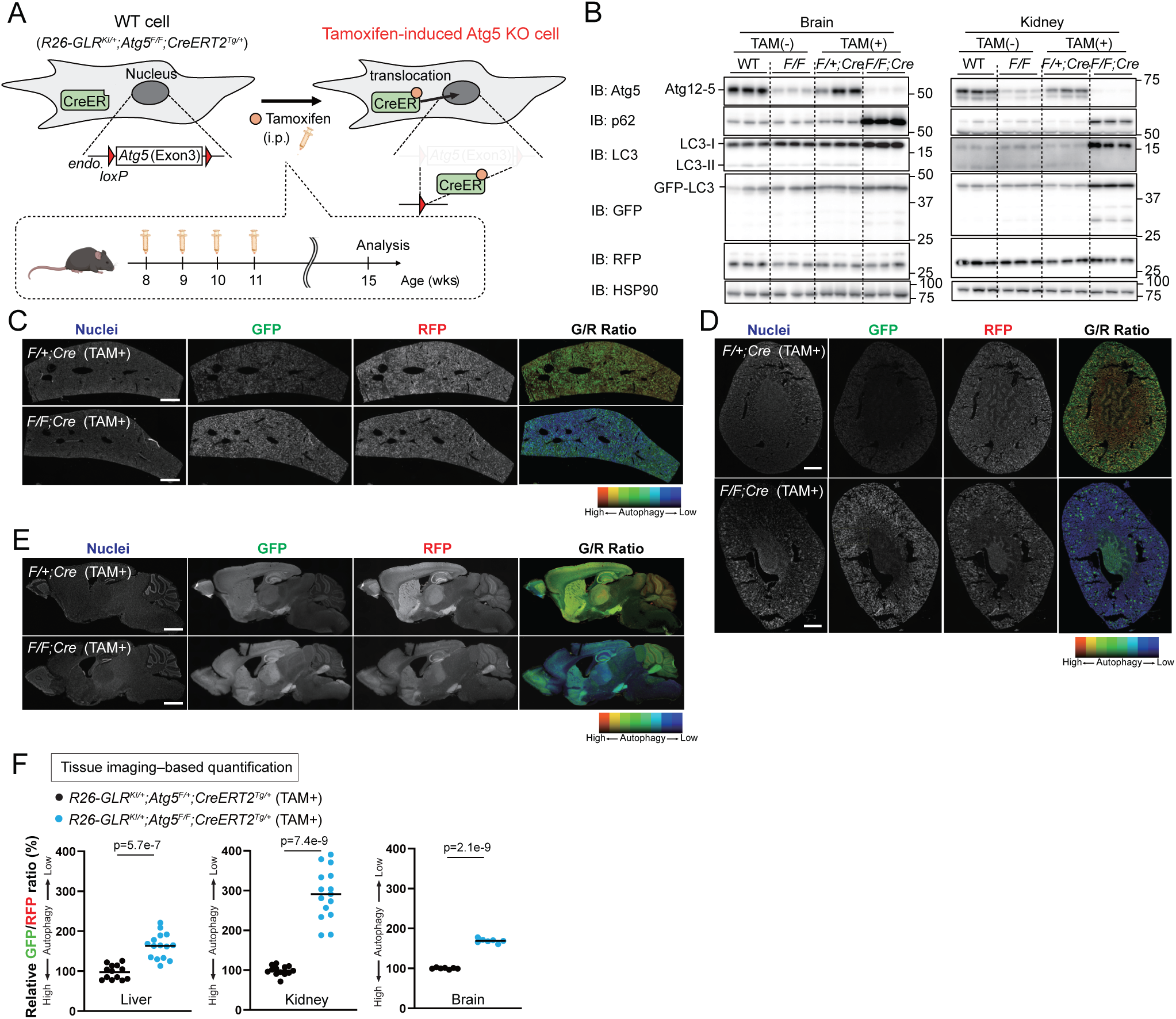
Distinct levels of basal autophagic flux between adult tissues. (A) Experimental design for tamoxifen-inducible Atg5 KO mice. Mice (3–4-month-old) were administered tamoxifen once per week for four weeks. One month after the last tamoxifen injection, mice were sacrificed for analysis. (B) Representative immunoblotting of the brain and kidney from *R26-GLR^KI/+^;Atg5^+/+^*, *R26-GLR^KI/+^;Atg5^flox/flox^*, *R26-GLR^KI/+^;Atg5^flox/+^;CreERT2^Tg/+^*, and *R26-GLR^KI/+^;Atg5^flox/flox^;CreERT2^Tg/+^* mice with or without tamoxifen administration. (C-E) Representative fluorescence and corresponding GFP/RFP ratio images of the liver (C), kidney (D), and brain (E) of *R26-GLR^KI/+^;Atg5^F/+^*;*CreERT2^Tg/+^*and *R26-GLR^KI/+^;Atg5^F/F^*;*CreERT2^Tg/+^*mice. Scale bar, 1 mm. (G) The GFP/RFP ratio calculated from tissue imaging of the liver, kidney, and brain of *R26-GLR^KI/+^;Atg5^F/+^*;*CreERT2^Tg/+^*and *R26-GLR^KI/+^;Atg5^F/F^*;*CreERT2^Tg/+^*mice. Each dot represents an individual mouse; bars indicate mean values; *n* = 7–15 per group. Differences between groups were analyzed by Welch’s *t*-test.

We then compared the GFP/RFP ratio between tamoxifen-treated *Atg5^F/+^;Cre* and *Atg5^F/F^;Cre* mice, representing autophagy-competent and autophagy-deficient conditions, respectively. Imaging of tissue sections revealed elevated GFP/RFP ratios in the liver (Figure 4C, F), kidney (Figure 4D, F), and brain (Figure 4E, F) of *Atg5^F/F^;Cre* mice compared with *Atg5^F/+^;Cre* mice, reflecting the accumulation of GFP signals caused by impaired autophagic degradation. These results suggest that, in contrast to starvation-induced autophagy, basal autophagy is active in the brain and may be comparable in magnitude to that in the liver.

To comprehensively assess basal autophagic flux across tissues, we quantified GFP/RFP ratios using the microplate-based assay. Each tissue displayed distinct GFP/RFP ratios, ranging from 130% to 350%, indicating marked tissue-specific differences in basal autophagic activity (Figure 5A). Moreover, no statistically significant differences were observed between male and female mice in any of the tissues examined, confirming that these tissue-specific patterns are consistent across both sexes (Figure 5—figure supplement 1). Pairwise statistical comparisons using the brain as a central reference revealed distinct functional groupings across tissues (Figure 5A and Supplementary Table 2). The kidney exhibited the highest basal autophagic flux, significantly exceeding all other tissues. The liver, pancreas, and lung displayed high autophagic flux comparable to the brain, showing no significant differences. Despite incomplete ATG5 deletion, the pancreas demonstrated a robust increase in autophagic flux, indicating substantial basal autophagic flux. In contrast, the small intestine, skeletal muscle, and heart exhibited significantly lower basal autophagic flux compared with the brain. Moreover, the increase in the GFP/RFP ratio upon Atg5 deletion was consistently greater in adult tissues than in embryonic tissues, indicating that basal autophagic flux becomes markedly elevated after birth (Figure 5B). Collectively, our results based on R26-GLR mice reveal pronounced stage- and tissue-specific differences in basal autophagic flux, which is generally low in embryos and differentially enhanced in adult tissues, including the brain.

**Figure 5.**
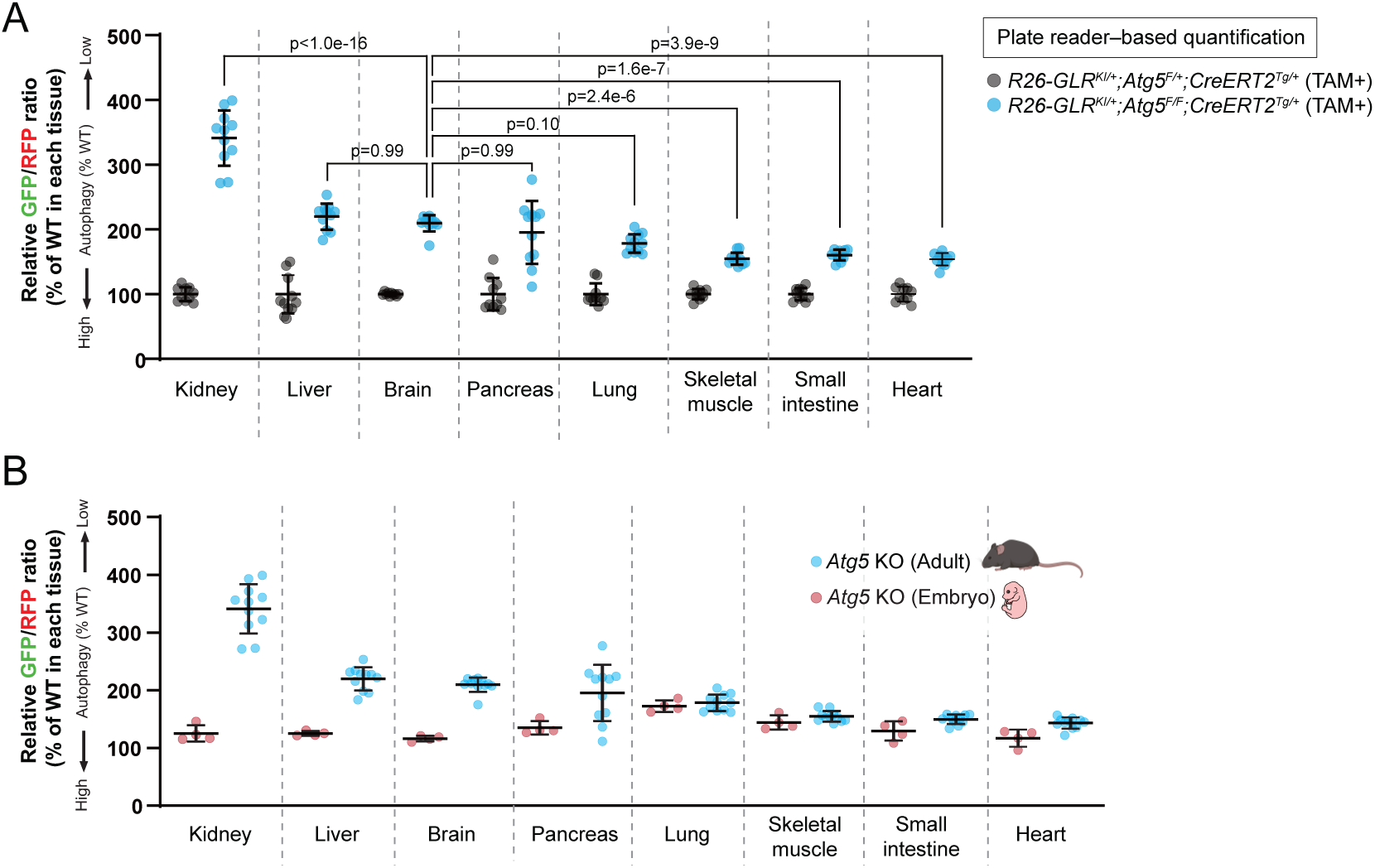
Distinct levels of basal autophagic flux in adult tissues and embryos. (A) Tissue-specific GFP/RFP ratios, as measured by microplate reader, of *R26-GLR^KI/+^;Atg5^F/F^*;*CreERT2^Tg/+^*mice, normalized to corresponding values of *R26-GLR^KI/+^;Atg5^F/+^*;*CreERT2^Tg/+^* mice (set as 100%). Each dot represents an individual mouse; bars indicate mean ± SD values; *n* = 10–11 per group. Differences across tissues and genotypes were analyzed by two-way ANOVA with Tukey’s post hoc test. (B) The GFP/RFP ratios of various tissues from Atg5 KO embryos and adult mice, normalized to the corresponding values in Atg5 WT controls (set as 100%). The data of embryos and adults are from Fig. 3E and Fig. 5A, respectively.

Given that basal autophagic activity is relatively high in the brain, we dissected regional differences within the brain by analyzing the GFP/RFP ratios in the cortex, hippocampus, and cerebellum. In *Atg5^F/+^;Cre* mice, the cortex and hippocampus showed comparable basal GFP/RFP ratios, whereas the cerebellum exhibited lower ratios (Figure 6—figure supplement 1A). However, comparison of these values with those of *Atg5^F/F^;Cre* mice revealed similar fold increases in these three regions (i.e., 1.5–1.6-fold increases), although the cerebellum might show a slightly lower increase (Figure 6—figure supplement 1B). To further refine this analysis at the cellular level, we quantified the GFP/RFP ratios in individual neurons. Specifically, we analyzed NeuN-positive neurons in the cortex (Figure 6A) and hippocampus (Figure 6B). Within the cerebellum, we focused on Purkinje cells (Figure 6C) due to their well-established vulnerability to autophagy deficiency, which leads to severe, cell-autonomous neurodegeneration (Hara et al., 2006; Komatsu et al., 2006; Komatsu et al., 2007; Liang et al., 2010). Consistent with the regional tissue data, all three evaluated neuronal types in Atg5F/F;Cre mice exhibited comparable increases in the GFP/RFP ratio relative to those in Atg5F/+;Cre mice (Figure 6D). These results suggest that basal autophagic flux is broadly comparable across brain regions and among the specific neuronal populations examined here.

**Figure 6.**
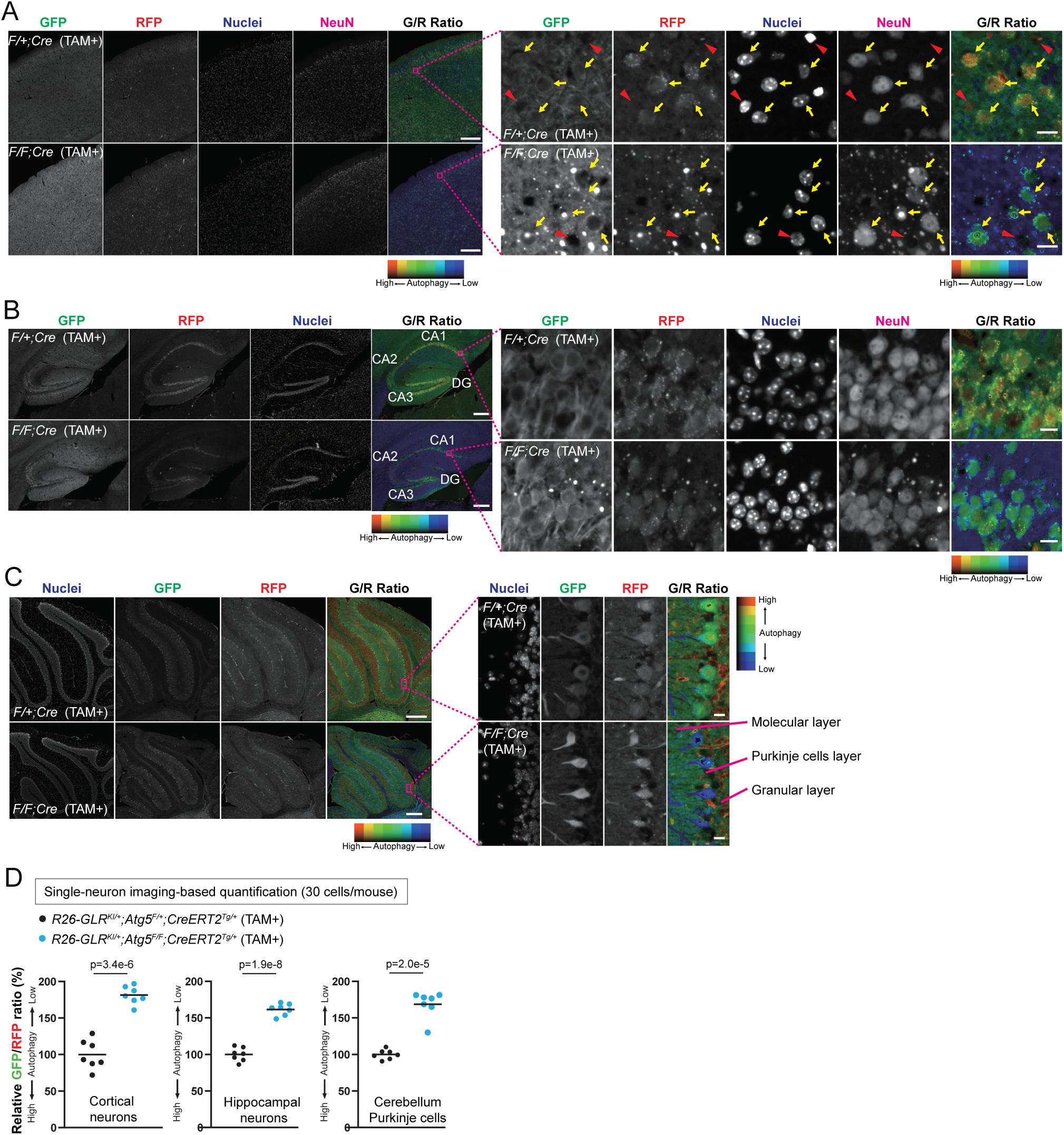
Comparable levels of basal autophagic flux among distinct neuronal populations. (A–C) Representative fluorescence and corresponding GFP/RFP ratio images of the cerebral cortex (A), hippocampus (B), and cerebellum (C) of *R26-GLR^KI/+^;Atg5^F/+^*;*CreERT2^Tg/+^*and *R26-GLR^KI/+^;Atg5^F/F^*;*CreERT2^Tg/+^*mice. Yellow arrows indicate Hoechst^+^/NeuN^+^ neurons, while red arrowheads indicate Hoechst^+/^NeuN^-^non-neuronal cells (B). Scale bars, 500 µm (low magnification) and 20 µm (high magnification). CA, Cornu Ammonis; DG, dentate gyrus. (D) The GFP/RFP ratio of neurons in the cortex and hippocampus and cerebellar Purkinje cells of *R26-GLR^KI/+^;Atg5^F/+^*;*CreERT2^Tg/+^*and *R26-GLR^KI/+^;Atg5^F/F^*;*CreERT2^Tg/+^*mice. Each dot represents an individual mouse (30 neurons were quantified per brain region for each mouse); bars indicate mean values; *n* = 7 per group. Differences between groups were analyzed by Welch’s *t*-test.

## Discussion

In this study, we developed a novel autophagy flux reporter mouse model, R26-GLR, and performed a systemic quantitative analysis of R26-GLR embryos and adult mice using tissue imaging and a plate reader-based method. One of the key findings is the unexpectedly high basal autophagic flux observed in the adult brain. Although a previous report indicated high basal autophagic flux in the liver through analysis of LC3 turnover with leupeptin (Haspel et al., 2011), quantitative assessment of the brain has been limited by the poor penetration of leupeptin across the blood–brain barrier. Neurons possess a highly polarized morphology, with autophagosomes forming at the distal axons and undergoing long-range retrograde transport for degradation in the soma (Maday & Holzbaur, 2014; Maday et al., 2012). This spatial uncoupling of autophagosome formation and degradation has long been inferred to mean that neuronal autophagy might be inefficient. Paradoxically, a previous study has reported the marked accumulation of autophagic vesicles in neurons following lysosomal inhibition (Boland et al., 2008), raising the possibility that neuronal autophagy is indeed robust. Our study provides, for the first time to our knowledge, direct *in vivo* evidence that basal autophagic flux is high in the brain, as well as the liver and kidney.

This high basal flux in the brain likely reflects the unique demands of neurons, which are long-lived, postmitotic cells that are highly vulnerable to the cumulative burden of proteotoxic and oxidative stress. Unlike dividing cells that can dilute damaged components through cell division, neurons must continuously maintain their proteome and organelle quality over a lifetime. Autophagy plays a pivotal role in this process by mediating the constitutive turnover of mitochondria, ER, and other subcellular components particularly prone to damage (Griffey & Yamamoto, 2022; Nixon & Rubinsztein, 2024). Furthermore, the complex architecture of neurons—spanning long axons, extensive dendrites, and numerous synaptic connections—necessitates localized regulation of protein and organelle turnover. Recent studies have begun to reveal that autophagy contributes to localized degradation in these subcellular compartments, particularly at presynaptic terminals and dendritic spines, where neuronal activity induces dynamic remodeling (Kallergi et al., 2022; Keary et al., 2023; Kulkarni et al., 2021; Shen et al., 2020; Soukup et al., 2016). This spatially restricted activity likely supports sustained basal flux by enabling local turnover in response to neuronal activity. Moreover, the restricted entry of peripheral insulin into the brain as a consequence of the blood–brain barrier might dampen mTORC1 activity and thereby promote autophagy. Although the molecular mechanisms remain incompletely defined, transcription factors such as TFEB, a key mediator of activity-dependent regulation of lysosomal biogenesis and autophagy, might help couple neuronal activity with global autophagic control (Agostini et al., 2024; Wang et al., 2023). Notably, such global mechanisms might cooperate with local activity-dependent regulation to establish robust, multi-layered control of autophagy in neurons. These insights raise the possibility that neuronal activity actively sustains autophagic flux to preserve synaptic and cellular homeostasis throughout life. Furthermore, this high baseline flux provides critical context for understanding the brain’s unique response to nutrient deprivation. While previous studies using tandem fluorescent LC3 (tfLC3) reporters noted region-specific activation of neural autophagy upon starvation (Alirezaei et al., 2010; Kaushik et al., 2011; Oliva Trejo et al., 2020), our findings confirm that the adult brain does not undergo the massive, organ-wide flux induction observed in metabolic tissues such as the liver and muscle (Mizushima et al., 2004). Consistent with our immunoblotting data (Figure 1—figure supplement 1F) and previous studies demonstrating the starvation resistance of neural autophagy (Kaade et al., 2024; Mizushima et al., 2004), neural mTORC1 activity remains exceptionally resistant to systemic nutrient deprivation.

A second major insight of the present study is the dynamic regulation of autophagy across development, highlighting how physiological demand shapes flux over time. Basal autophagic flux was uniformly low during embryogenesis but increased markedly after birth, likely reflecting the shift from a stable fetal environment to a more variable postnatal environment (Kuma et al., 2004). Prior studies using mitophagy and ER-phagy reporter mice have also demonstrated increases in selective autophagy throughout development (McWilliams et al., 2016; Sang et al., 2024), supporting the notion that autophagic demand rises in mature adult tissues compared to embryonic stages. Notably, this increase was highly tissue-specific. This non-uniformity highlights the distinct physiological reliance of individual organs on constitutive autophagy postnatally: the kidney (handling continuous filtration and reabsorption), the liver (serving as a central metabolic hub), the brain, and the pancreas (synthesizing and secreting massive amounts of digestive enzymes) operate under constant metabolic and proteotoxic strain. In contrast, other tissues, such as skeletal muscle and small intestine, maintained relatively low baseline flux. Particularly in the neural system, transient and localized autophagy during early development plays important roles in processes such as axonal development (Lv et al., 2014) and microglial synaptic pruning (Kim et al., 2017; Lieberman et al., 2019). Nevertheless, mature postmitotic cells—particularly neurons—cannot dilute misfolded or aggregated proteins or damaged organelles through cell division. Consequently, accommodating postnatal environmental stimuli while preserving lifelong tissue homeostasis is thought to drive a continuously higher demand for baseline autophagic flux in adult postmitotic tissues compared to embryonic stages. This spatial and temporal specificity might help explain the severe, tissue-specific phenotypes that emerge postnatally in autophagy-deficient mice, particularly in the brain (Hara et al., 2006; Komatsu et al., 2006; Komatsu et al., 2007; Liang et al., 2010), liver(Komatsu et al., 2005; Takamura et al., 2011; Yoshii et al., 2016), and kidney (Kawakami et al., 2015). These insights have direct implications for human disease; mutations in core autophagy genes frequently cause neurodevelopmental and neurodegenerative disorders (Almannai et al., 2022; Collier et al., 2021; Haack et al., 2012; Jelani et al., 2019; Kim et al., 2016; Maroofian et al., 2021; Saitsu et al., 2013; Suleiman et al., 2018), likely via impaired homeostasis in neural tissues that rely heavily on constitutive autophagy.

In conclusion, we established the versatile R26-GLR mouse model, which enables both single-cell-level analysis of tissue imaging data and semi-high-throughput quantification of autophagic flux using a microplate reader-based method. Using this mouse model, our study elucidated a spatiotemporal landscape of basal autophagy, revealing distinct organ- and stage-specific patterns, with particularly high basal autophagic flux in the brain. Although conducted primarily at the tissue level, our analysis lays the foundation for future cell-type–specific investigations, particularly in complex organs such as the brain. In addition, incorporating other Atg-deficient mouse models will further validate and extend these findings. Together, these efforts will provide a framework for understanding the physiological roles of autophagy as well as the consequences of its disruption in disease contexts.

## Limitations of this study

Although the R26-GLR mouse model is a versatile tool for *in vivo* assessment of autophagic flux, several limitations should be considered. GFP–LC3 and RFP are both subject to degradation not only via autophagy but also through non-autophagic pathways, particularly the ubiquitin–proteasome system. The relative contribution of these degradation pathways is assumed to vary among tissues and cell types. Therefore, differences in GFP/RFP ratios between tissues or cell types should not be interpreted as absolute levels of autophagic flux. To determine whether changes in the reporter reflect true autophagy-dependent processes, we recommend parallel analyses under autophagy-deficient conditions. In addition, enzymes, such as UBA6 and BIRC6, which mediate proteasomal degradation of LC3-I (Jia & Bonifacino, 2019; Jiang et al., 2019), might differentially affect GFP–LC3 stability and thereby influence the reporter signal, particularly in pathological states like cancer, in which BIRC6 has been reported to be upregulated (Hu et al., 2015; Low et al., 2013; Ren et al., 2005). Finally, within the neural system, our single-neuron quantification was restricted to specific, visually demarcated neuronal populations (such as cortical/hippocampal principal neurons and Purkinje cells), leaving a systematic survey across the full diversity of neuronal subtypes (e.g., excitatory vs. inhibitory neurons across all regions) unaddressed. Future studies combining cell-type-specific Cre drivers or cell-isolation techniques with the R26-GLR reporter will be essential to comprehensively map subtype-specific autophagic flux across diverse neuronal and glial populations.

## Materials and methods

### Key resources table

### Materials availability

This study generated novel mouse lines, which have been deposited at the RIKEN BioResource Research Center (BRC) and are available for distribution: R26-STOP-GLR (RBRC12610) and R26-GLR (RBRC12611). Requests for materials should be directed to the BRC.

### Mouse Strains and husbandry

All animal experiments were approved by the Institutional Animal Care and Use Committee of the University of Tokyo (A2023M010-03). Mice were housed, two to seven individuals per cage, at 20–21℃ in a humidity-controlled room and maintained on a 12 h light/dark cycle (08:00 to 20:00 light on), with a normal chow diet (CE-2, CLEA), and water was provided ad libitum. Cages and bedding were changed once per week. The following mouse strains were used and maintained on a C57BL/6J background by crossing with wild-type C57BL/6J mice (Japan SLC): *Atg5^+/-^* (Kuma et al., 2004), *Atg5^flox/+^* (Hara et al., 2006), *Vasa-Cre^Tg/+^* (Gallardo et al., 2007), *Ubc-CreERT2^Tg/+^*(Ruzankina et al., 2007), R26-STOP-GLR (in this study), and R26-GLR (in this study).

### Animals and procedures

For starvation treatment, 3- to 4-month-old R26-GLR mice of either sex were starved for 24 or 48 h under free access to drinking water. For the *Atg5* KO embryo experiment, R26-GLR mice were crossed with *Atg5^+/-^* mice to prepare *R26-GLR^KI/+^;Atg5^+/+^* and *R26-GLR^KI/+^;Atg5^-/-^*mice of either sex. Embryos were collected at embryonic day 19.5 by caesarean section. For the tamoxifen-induced *Atg5* KO experiment, R26-GLR mice were crossed with *Atg5^flox/+^* and *Ubc-CreERT2^Tg/+^* mice to prepare *R26-GLR^KI/+^;Atg5^flox/+^;Ubc-CreERT2^Tg/+^*and *R26-GLR^KI/+^;Atg5^flox/flox^;Ubc-CreERT2^Tg/+^*mice of either sex. Eight- to nine-week-old mice were intraperitoneally administrated tamoxifen once per week for 4 weeks as described previously (Yang et al., 2022). In brief, tamoxifen was dissolved at 20 mg/mL concentration in a mixture of 98% sunflower seed oil and 2% ethanol, and a dose of 150 µL per 25 g body weight was injected at the indicated time. One month following the last tamoxifen administration, mice were sacrificed. Mice were sacrificed under anesthesia with isoflurane (brief exposure for ∼1 min to induce deep anesthesia) at the indicated time points in each experiment, and tissues were dissected immediately after PBS perfusion. For biochemical analyses, tissues were snap-frozen in liquid nitrogen and stored at −80℃ until use. For tissue imaging to analyze the GFP/RFP ratio, tissues were fixed in 4% paraformaldehyde (PFA) in phosphate-buffered saline (PBS) overnight. Fixed tissues were subjected to 15% sucrose and 30% sucrose in PBS until they were submerged and embedded in Tissue-Tek OCT Compound (Sakura Finetek Japan), stored at −80℃ until use, and finally sectioned using a cryostat (Leica CM1950) to a 7-µm thickness.

### Generation of R26-STOP-GLR knock-in mouse

The targeting vector contains a splice CAG promoter, frt-flanked repeating SV40 polyadenylation signal (pA), STOP cassette containing the terminator of the yeast His3 gene, SV40 pA signal, and GFP-LC3-RFP-WPRE-bGHpA. This vector has 5′ and 3′ homology arms of 4.7 and 5.2 kbp, respectively, which target the construct to the Xba1 site of intron 1 at the Rosa26 locus. The targeting vector was linearized and electroporated into the RENKA C57BL/6 embryonic stem cell line(Mishina & Sakimura, 2007). G418-resistant ES clones were screened by southern blot analysis for homologous recombination at the Rosa26 locus by probing SpeI-digested genomic DNA with a 0.4-kb genomic fragment from immediately upstream of the 5′ arm and EcoRI-digested genomic DNA with a 0.6-kb genomic fragment from immediately downstream of the 3′ arm. Targeted ES clones were injected into eight-cell stage ICR, which were cultured to produce blastocysts and later transferred to pseudopregnant ICR females. The resulting male chimeric mice were crossed with female C57BL/6 mice to establish the R26-STOP-GLR line.

### Genotyping

For the determination of mice genotype, approximately 1-mm-long tail samples were mixed with 100 µL of Tail Lysis buffer (50 mM Tris-HCl (pH 8.0), 2 mM NaCl, 1 mM EDTA (pH 8.0), 0.063% SDS) and 1 µL of Proteinase K (Nacalai Tesque Cat#15679-06), heated at 55℃ overnight, and then finally boiled at 98°C for 1 min to inactivate Proteinase K. After centrifugation at room temperature at 13,500 × *g* for 10 min, supernatants were used for PCR. PCR was performed using Quicktaq (TOYOBO #DTM-101) with the following thermal cycling program: 94℃ for 2 min; 35 cycles of 94℃ for 30 s, 55℃ for 30 s, and 68℃ for 2 min; and a final extension at 68℃ for 1 min. PCR products were electrophoresed in 2% agarose gel containing MIDORI Green Advance (NIPPON Genetics #NE-MG04). The primers used in this study are listed (Supplementary Table 1).

### Immunoblotting

Lysis buffer containing 1% Triton X-100 in PBS or RIPA buffer containing 50 mM Tris-HCl (pH 8.0), 150 mM NaCl, 0.5% (*w*/*v*) sodium deoxycholate, 0.1% (*w*/*v*) SDS, and 1.0% (*w*/*v*) NP-40 substitute (FUJIFILM Wako Cat#182-02451), supplemented with complete EDTA-free protease inhibitor cocktail (Roche Cat#11836170001) was added to the tissues and homogenized by Polytron Homogenizer (KINEMATICA pT-MR2500E) or multi-beads Shocker (Yasui Kikai MB3200C(S)). The supernatants were collected by centrifugation at 2,500 × *g* for 10 min, and the protein concentrations were adjusted using the Bradford method. The tissue lysates were solubilized with immunoblot sample buffer (46.7 mM Tris-HCl (pH 6.8), 1.67% sodium dodecyl sulfate, 5% glycerol, 1.55% dithiothreitol, and 0.003% bromophenol blue). Twenty-microgram total protein samples were subjected to SDS-polyacrylamide gel electrophoresis, transferred to an Immobilon-P polyvinylidene difluoride membrane (Millipore IPVH00010), and blotted with antibodies. Super-Signal West Pico Chemiluminescent substrate (Thermo Fisher Scientific Cat#34580) or Immobilon Western Chemiluminescent HRP Substrate (Merck Millipore Cat#WBKLS0500) was used for detection of each protein signal. Signals were captured using the FUSION SOLO7S instrument (Vilber-Lourmat). The resulting images were processed using Image J software.

### Antibodies

For immunoblotting, the following primary antibodies were used at a 1:1,000 dilution: Anti-LC3 (NM1) (Quy et al., 2013), anti-LC3B (Novus Biologic Cat#NB100-2220), anti-Lamin A/C (Cell Signaling Technology Cat#2032), anti-GFP (Invitrogen Cat#A6455), anti-RFP (MBL Cat#M208-3), anti-HSP90 (BD Biosciences Cat#610419), anti-ATG5 (Sigma-Aldrich Cat#A0731), anti-pS6K1(T389) (Cell Signaling Technology Cat#9234), anti-4E-BP1 (Cell Signaling Technology Cat#9452), and anti-p62 (MBL Cat#PM045) antibodies. For immunofluorescence, the following primary antibodies were used at a 1:200 dilution: Anti-NeuN (Abcam Cat# ab177487).

Secondary antibodies used for immunoblotting were horseradish peroxidase-conjugated anti-mouse IgG (Jackson ImmunoResearch Laboratories Cat#315-036-003) and anti-rabbit IgG (Jackson ImmunoResearch Laboratories Cat#111-035-144) at a 1:10,000 dilution. For immunofluorescence, Alexa Fluor-conjugated secondary antibodies (Invitrogen Cat#A-21245) were used at a 1:500 dilution.

### Microscopic observations

Cryosections were washed with PBS, stained with Hoechst33342 (Immunochemistry Technologies Cat#639) in PBS for 10 min, and then washed three more times with PBS. When immunofluorescence was required, sections were first permeabilized with 0.1% Triton X-100 in PBS for 15 min at room temperature, followed by blocking with 5% BSA. The sections were then incubated with primary antibodies overnight at 4°C. After washing, sections were incubated with appropriate secondary antibodies for 1 hour at room temperature, followed by Hoechst 33342 staining. The coverslips were mounted with SlowFade antifade reagent. Prepared cryosections were observed using an all-in-one fluorescence microscope (Keyence BZ-X800) equipped with 4×, 20×, and 60× objective lenses and a fluorescence filter cube for DAPI (Keyence OP-87762), GFP (Keyence OP-87763), TexasRed (Keyence OP-87765), and Cy5 (Keyence OP-87766) or using a confocal microscope (Olympus FV1000) equipped with 10× and 60× objective lenses (Olympus UPLXAPO10X and UPLXAPO60XO). Images were captured with BZ-X analyzer software (Keyence) or FluoView software (Olympus FV10-ASW) and processed using Image J software (NIH).

### Measurement of the GFP/RFP ratio by tissue fluorescent imaging

All imaging and quantitative analyses were performed in a blinded manner. Specifically, after sample information was blinded by a third party, fluorescence imaging and analysis were conducted using MetaMorph software (Molecular Devices). For the starvation-treated *Atg5* KO embryo and *Atg5* KO adult experiments, one tissue section from each mouse was used, and the whole tissue areas were imaged and analyzed. For the *Atg5* KO embryo experiment, one tissue section from each mouse was used and 5–10 areas of each tissue were imaged and analyzed. For single-neuron assessment in the *Atg5* KO adult brain, 30 neurons of each type were analyzed per mouse. Cortical and hippocampal neurons were identified by NeuN staining; the faint cytoplasmic signal, in addition to nuclear staining, allowed for manual outlining of the cell bodies. For Purkinje cells, cell bodies were identified and manually outlined based on the RFP signal. In brief, gray-scale ratio images were generated by the “Arithmetic” tool in MetaMorph software using the observed GFP and RFP channel images. The GFP/RFP ratio was calculated by averaging the GFP/RFP ratio of each pixel in the selected tissue area. To generate the GFP/RFP ratio images with pseudo color ranging from blue to red, background signals in tissues-free areas were subtracted from RFP channel images. Then, using the “Ratio Images” tool in MetaMorph, GFP/RFP ratio images were generated. The dynamic range of the GFP/RFP ratio images was determined by the “Min Ratio” and “Max Ratio” scores, which were unified among the samples of each experiment.

### Quantification and normalization of relative autophagic flux

To evaluate autophagic flux across tissues, all mice and embryos analyzed carried the reporter allele (*R26-GLR^KI/+^*). The raw GFP/RFP ratio of each individual sample (*Atg5* KO: adult *Atg5^F/F^;Cre* [*R26-GLR^KI/+^;Atg5^flox/flox^;Ubc-CreERT2^Tg/+^*] or embryonic *Atg5^Δ/Δ^* [*R26-GLR^KI/+^;Atg5^-/-^*]) was normalized to the mean raw GFP/RFP ratio of the corresponding control group (*Atg5^F/+^;Cre* [*R26-GLR^KI/+^;Atg5^flox/+^;Ubc-CreERT2^Tg/+^*] for adults; *Atg5^+/+^* [*R26-GLR^KI/+^;Atg5^+/+^*] for embryos) for that specific tissue, with the control group mean defined as 100%.

### Microplate reader measurement of the GFP/RFP ratio

Tissue homogenates were prepared in the same method described for immunoblotting. In brief, for each sample, nine times the volume of lysis buffer (µL) to the tissue weights (mg) were added and homogenized. After centrifugation, the soluble fractions were used as tissue homogenates. Fifty-microliter tissue lysates were mixed with 150 µL of PBS and loaded into a 96-well plate. Fluorescent signals of GFP and RFP were measured with a microplate reader (Enspire Multimode Plate Reader, PerkinElmer) to calculate the GFP/RFP ratio.

### Statistical analysis

All statistical analyses were performed using GraphPad Prism (GraphPad Software). For comparisons between two groups, Welch’s *t*-test (two-tailed) was applied. For multiple group comparisons, one-way analysis of variance (ANOVA) followed by Tukey’s multiple comparison test was used.

## Acknowledgments

We thank Prof. Toshiya Endo of Kyoto Sangyo University for the generous gift of the anti-RFP antibody used in the initial phase of this study. This work was supported by Exploratory Research for Advanced Technology (ERATO) research funding program of the Japan Science and Technology Agency (JST) (JPMJER1702 to N.M.), Grant-in-Aid for Specially Promoted Research (22H04919 to N.M.), Japan Society for the Promotion of Science (JSPS) KAKENHI (AdAMS) (Aa160004 to N.M.). This work was also supported by a PhD fellowship from the World-leading Innovative Graduate Study Program for Life Science and Technology (WINGS-LST), funded by the Ministry of Education, Culture, Sports, Science and Technology (MEXT), Japan (to Y.K.). The graphical abstract and figure schematics were created with BioRender.com.

## Author contributions

Conception and design, H.M. and N.M.; generation of reporter mice, H.M., Y.H., M.A., and K.S.; acquisition of data, Y.K.; interpretation of data, Y.K., T.E., H.M., N.M.; writing of the manuscript, Y.K., T.E., N.M.

## Declaration of AI-assisted technologies in the writing process

The authors used ChatGPT (GPT-4.0) to assist with language refinement during manuscript preparation. The final content was thoroughly reviewed and edited by the authors, who take full responsibility for the publication.

**Figure 1—figure supplement 1. The generation and validation of R26-GLR mice and their starvation response.** (A) Schematic representation of the R26-STOP-GLR and R26-GLR alleles. The positions of primers used for genotyping are indicated (left). Genotypes tested (gray box) and corresponding PCR results (right) are shown. Primer sets #1–6, along with internal control (I.C.) primers, were used. M, DNA ladder. (B) Representative immunoblotting from WT (W), R26-STOP-GLR (S), and R26-GLR (G) mice (3–4-month-old). (C) Representative tissue fluorescence images of WT and R26-GLR mice (3–4-month-old) obtained using 10× and 60× objective lenses. Scale bars, 500 µm (low magnification) and 40 µm (high magnification). (D, E, F) Representative immunoblotting results of the liver (D), skeletal muscle (E), and brain (F) from mice starved for 24 or 48 h. Asterisks indicate non-specific bands. (G, H, I) Quantification of the GFP/RFP ratio determined by immunoblotting of tissue lysates from the liver (G), skeletal muscle (H), and brain (I) of mice starved for 24 or 48 h. Each dot represents an individual mouse; bars indicate mean values; *n* = 8–10 per group. Differences among groups were analyzed by one-way ANOVA with Tukey’s post hoc test.

**Figure1—figure supplement 2. GFP-LC3-RFP reporter expression in additional tissues of R26-GLR mice.** Representative tissue fluorescence images of the heart, lung, spleen, and small intestine from WT and R26-GLR mice (3–4-month-old) obtained using 10× and 60× objective lenses. In the spleen, WP and RP indicate white pulp and red pulp, respectively. Scale bars, 500 µm (low magnification) and 40 µm (high magnification).

**Figure1—figure supplement 3. Expression of the R26-GLR reporter does not substantially affect endogenous autophagic activity.** (A) Representative immunoblots in various tissues (brain, liver, kidney, spleen, pancreas, and skeletal muscle) from wild-type (WT) and *R26-GLR^KI/+^* mice (*n* = 3 per group; male and female as indicated). Blue asterisks indicate non-specific bands. (B) Quantification of p62 protein levels in WT and *R26-GLR^KI/+^* mice across tissues. Each dot represents an individual mouse; bars indicate mean values normalized to expression of HSP90; *n* = 3 per group. Differences between groups were analyzed by Welch’s t-test.

**Figure 2—figure supplement 1. Validation of the microplate reader–based GFP/RFP ratio measurement.** (A) Soluble fractions from kidney lysates of WT and R26-GLR mice (3–4-month-old) were serially diluted and measured using a microplate reader (*n* = 3 per group). Data are presented as mean ± SD values. The coefficients of determination (*R*²) from linear regression of GFP signal measurements and lysate concentration for R26-GLR mice is shown. (B) Schematic overview of the experimental conditions used for microplate reader–based detection. (C) Liver and kidney tissues from WT and R26-GLR mice (3–4-month-old) were homogenized in either 1% Triton X-100/PBS or RIPA buffer and then subjected to microplate reader analysis (*n* = 3 per group). Bars represent mean ± SD values. (D) Liver and kidney lysates from WT and R26-GLR mice (3–4-month-old) were centrifuged at different speeds and analyzed using a microplate reader (*n* = 3 per group). Bars represent mean ± SD values. (E) External appearance of the liver lysate prepared under different conditions. (F) Soluble and insoluble fractions from liver and kidney lysates of WT and R26-GLR mice (3–4-month-old) were subjected to microplate reader analysis (*n* = 3 per group). Bars represent mean ± SD values. (G) Representative immunoblotting results of soluble and insoluble fractions of the liver and kidney.

**Figure 3—figure supplement 1. Validation of R26-GLR reporter expression and Atg5 deletion in embryonic tissues.** (A) Representative immunoblotting of head and body tissues from Atg5 wild-type (WT), heterozygous (HT), or knockout (KO) embryos with or without the GFP–LC3–RFP reporter. (B) Soluble fractions from the head and body lysates of WT and R26-GLR embryos were serially diluted and measured using a microplate reader (*n* = 3 per group). Data are presented as mean ± SD values. The coefficients of determination (*R*²) of linear regressions of measurements of tissue concentration and GFP signal for R26-GLR embryos are shown. (C) Representative tissue fluorescence images of WT and R26-GLR embryos. Scale bars, 1 mm (body) and 200 µm (head). (D) Representative GFP/RFP ratio images of the head of *Atg5* WT and *Atg5* KO embryos. Scale bar, 200 µm. (E) Representative GFP/RFP ratio images of kidney glomeruli of *Atg5* WT and *Atg5* KO embryos. Yellow arrows indicate podocytes. Scale bar, 100 µm.

**Figure 4—figure supplement 1. Validation of tamoxifen-induced Atg5 knockout and dynamic differences in basal autophagy between embryonic and adult tissues.** (A) Schematic illustration of Atg5 WT, flox, and deleted alleles. The positions of the PCR primers are indicated. (B) Representative genomic PCR results of *R26-GLR^KI/+^;Atg5^flox/flox^*, *R26-GLR^KI/+^;Atg5^flox/+^;CreERT2^Tg/+^*, and *R26-GLR^KI/+^;Atg5^flox/flox^;CreERT2^Tg/+^* mice with or without tamoxifen administration. Tail samples were used for PCR. (C) Quantification of the ATG5 and p62 protein expression levels in *R26-GLR^KI/+^;Atg5^flox/flox^*, *R26-GLR^KI/+^;Atg5^flox/+^;CreERT2^Tg/+^*, and *R26-GLR^KI/+^;Atg5^flox/flox^;CreERT2^Tg/+^* mice with or without tamoxifen administration. Each dot represents an individual mouse; bars indicate mean values normalized to HSP90 expression (*n* = 3 per group). Differences between groups were analyzed by Welch’s *t*-test. (D) Representative immunoblotting of major tissues from *R26-GLR^KI/+^;Atg5^F/+^*;*CreERT2^Tg/+^* and *R26-GLR^KI/+^;Atg5^F/F^*;*CreERT2^Tg/+^* mice. (E) Quantification of the ATG5 expression levels in *R26-GLR^KI/+^;Atg5^F/+^*;*CreERT2^Tg/+^*and *R26-GLR^KI/+^;Atg5^F/F^*;*CreERT2^Tg/+^* mice. Each dot represents an individual mouse; bars indicate mean values normalized to expression of HSP90; *n* = 3 per group. Differences between groups were analyzed by Welch’s *t*-test. (F) Microplate reader-based measurement of the GFP/RFP ratio in the liver, kidney, and brain of *R26-GLR^KI/+^;Atg5^F/+^*;*CreERT2^Tg/+^* and *R26-GLR^KI/+^;Atg5^F/F^*;*CreERT2^Tg/+^* mice. Each dot represents an individual mouse; bars indicate mean values; *n* = 10–11 per group. Differences between groups were analyzed by Welch’s *t*-test.

**Figure5—figure supplement 1. Evaluation of potential sex differences in basal autophagic flux across tissues.**

Tissue-specific GFP/RFP ratios, as measured by microplate reader, of *R26-GLR^KI/+^;Atg5^F/F^*;*CreERT2^Tg/+^*mice, normalized to corresponding values of *R26-GLR^KI/+^;Atg5^F/+^*;*CreERT2^Tg/+^*mice (set as 100%), plotted separately for male and female mice (derived from the same dataset as in Figure 5A). Each dot represents an individual mouse; bars indicate mean values; *n* = 5–6 per group. Differences between sexes across tissues were analyzed by two-way ANOVA with Tukey’s post hoc test.

**Figure 6—figure supplement 1. Comparable levels of basal autophagic flux across brain regions.** (A) Region-specific GFP/RFP ratios (raw values) in the brain of *R26-GLR^KI/+^;Atg5^F/+^*;*CreERT2^Tg/+^* and *R26-GLR^KI/+^;Atg5^F/F^*;*CreERT2^Tg/+^* mice measured by tissue imaging. Each dot represents an individual mouse; bars indicate mean values; *n* = 7 per group. Differences among groups were analyzed by two-way ANOVA with Tukey’s post hoc test. (B) Region-specific GFP/RFP ratios (normalized values). The data shown in (A) were normalized to values in each corresponding region of *R26-GLR^KI/+^;Atg5^F/+^*;*CreERT2^Tg/+^*mice (set as 100%). Differences among groups were analyzed by two-way ANOVA with Tukey’s post hoc test.

