## Supplementary Figures for "High basal autophagic activity in the brain revealed by systemic quantitative analysis using GFP-LC3-RFP mice"

### Figure 1—figure supplement 1

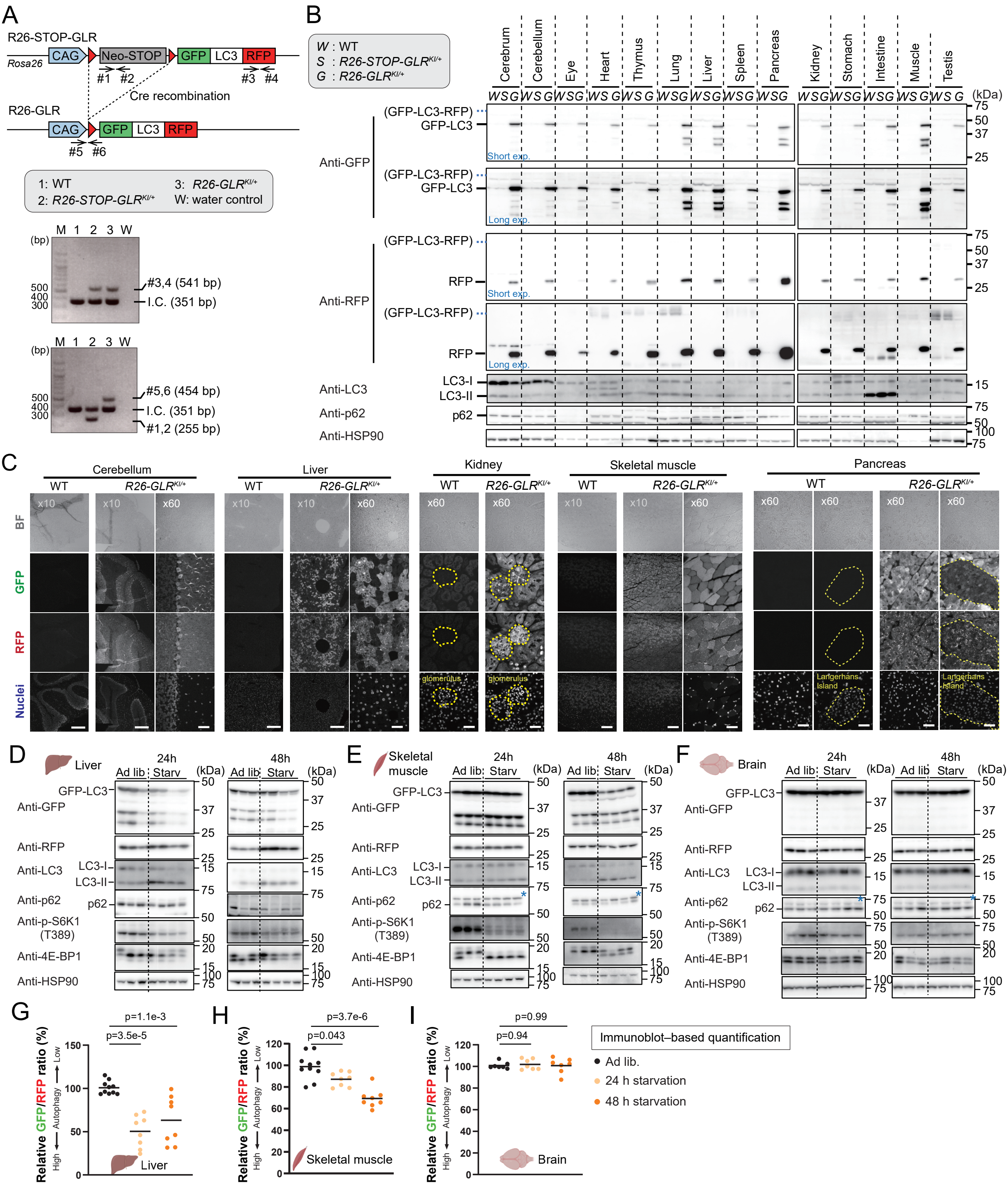

**Figure 1—figure supplement 1. The generation and validation of R26-GLR mice and their starvation response.**
(A) Schematic representation of the STOP-R26-GLR and R26-GLR alleles. The positions of primers used for genotyping are indicated (left). Genotypes tested (gray box) and corresponding PCR results (right) are shown. Primer sets #1–6, along with internal control (I.C.) primers, were used. M, DNA ladder.
(B) Representative immunoblotting from WT (W), STOP-R26-GLR (S), and R26-GLR (G) mice (3–4-month-old).
(C) Representative tissue fluorescence images of WT and R26-GLR mice (3–4-month-old) obtained using 10× and 60× objective lenses. Scale bars, 500 µm (low magnification) and 40 µm (high magnification).
(D, E, F) Representative immunoblotting results of the liver (D), skeletal muscle (E), and brain (F) from mice starved for 24 or 48 h. Asterisks indicate non-specific bands.
(G, H, I) Quantification of the GFP/RFP ratio determined by immunoblotting of tissue lysates from the liver (G), skeletal muscle (H), and brain (I) of mice starved for 24 or 48 h. Each dot represents an individual mouse; bars indicate mean values; $n$ = 8–10 per group. Differences among groups were analyzed by one-way ANOVA with Tukey's post hoc test.

### Figure 1—figure supplement 2

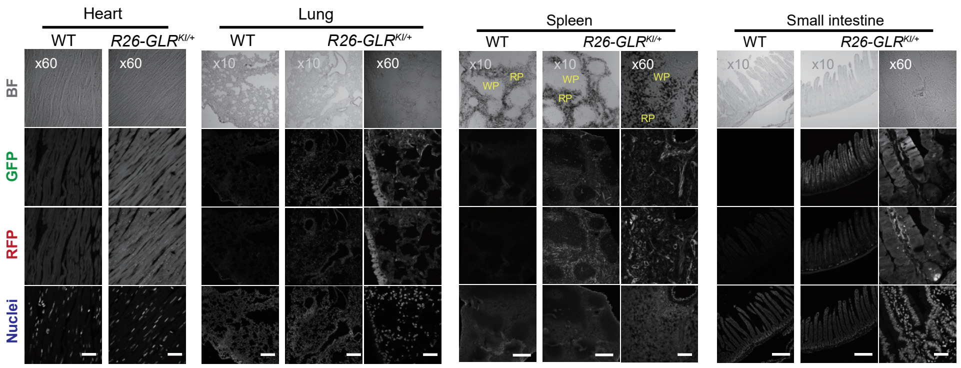


**Figure1—figure supplement 2. GFP-LC3-RFP reporter expression in additional tissues of R26-GLR mice.** Representative tissue fluorescence images of the heart, lung, spleen, and small intestine from WT and R26-GLR mice (3–4-month-old) obtained using 10× and 60× objective lenses. In the spleen, WP and RP indicate white pulp and red pulp, respectively. Scale bars, 500 µm (low magnification) and 40 µm (high magnification).

### Figure 1—figure supplement 3

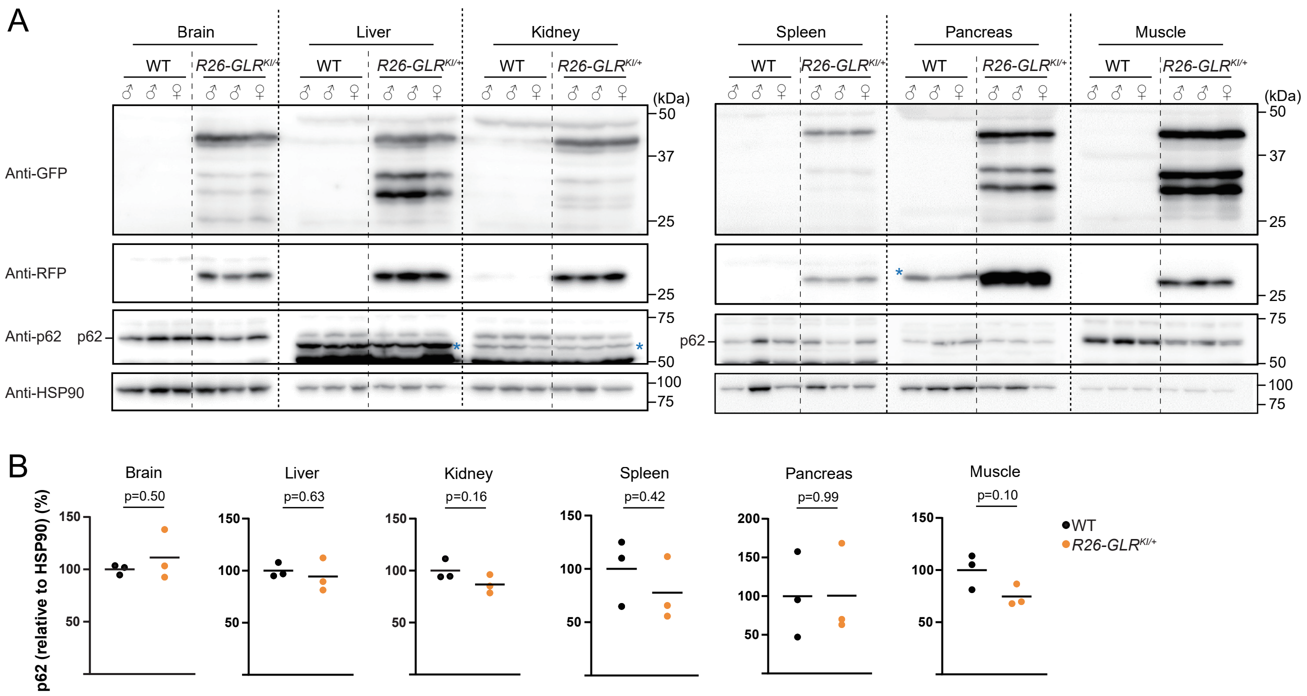


**Figure1—figure supplement 3. Expression of the R26-GLR reporter does not substantially affect endogenous autophagic activity.**

(A) Representative immunoblots in various tissues (brain, liver, kidney, spleen, pancreas, and skeletal muscle) from wild-type (WT) and *R26-GLR*$^{KI/+}$ mice ($n$ = 3 per group; male and female as indicated). Blue asterisks indicate non-specific bands.

(B) Quantification of p62 protein levels in WT and *R26-GLR*$^{KI/+}$ mice across tissues. Each dot represents an individual mouse; bars indicate mean values normalized to expression of HSP90; $n$ = 3 per group. Differences between groups were analyzed by Welch's t-test.

Figure 2—figure supplement 1

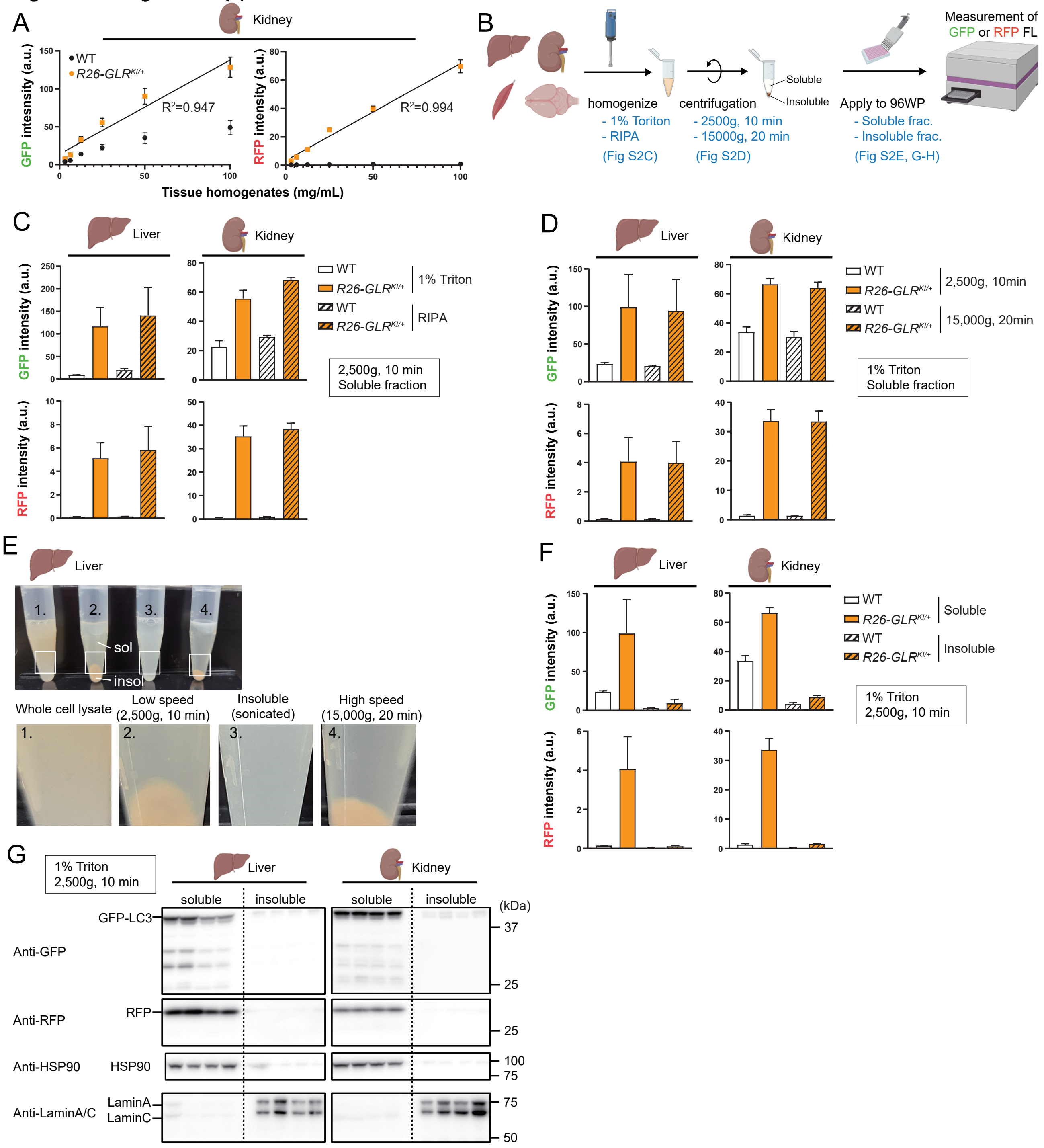

**Figure 2—figure supplement 1. Validation of the microplate reader–based GFP/RFP ratio measurement.**
(A) Soluble fractions from kidney lysates of WT and R26-GLR mice (3–4-month-old) were serially diluted and measured using a microplate reader (*n* = 3 per group). Data are presented as mean ± SD values. The coefficients of determination ($R^2$) from linear regression of GFP signal measurements and lysate concentration for R26-GLR mice is shown.
(B) Schematic overview of the experimental conditions used for microplate reader–based detection.
(C) Liver and kidney tissues from WT and R26-GLR mice (3–4-month-old) were homogenized in either 1% Triton X-100/PBS or RIPA buffer and then subjected to microplate reader analysis (*n* = 3 per group). Bars represent mean ± SD values.
(D) Liver and kidney lysates from WT and R26-GLR mice (3–4-month-old) were centrifuged at different speeds and analyzed using a microplate reader (*n* = 3 per group). Bars represent mean ± SD values.
(E) External appearance of the liver lysate prepared under different conditions.
(F) Soluble and insoluble fractions from liver and kidney lysates of WT and R26-GLR mice (3–4-month-old) were subjected to microplate reader analysis (*n* = 3 per group). Bars represent mean ± SD values.
(G) Representative immunoblotting results of soluble and insoluble fractions of the liver and kidney.

### Figure 3—figure supplement 1

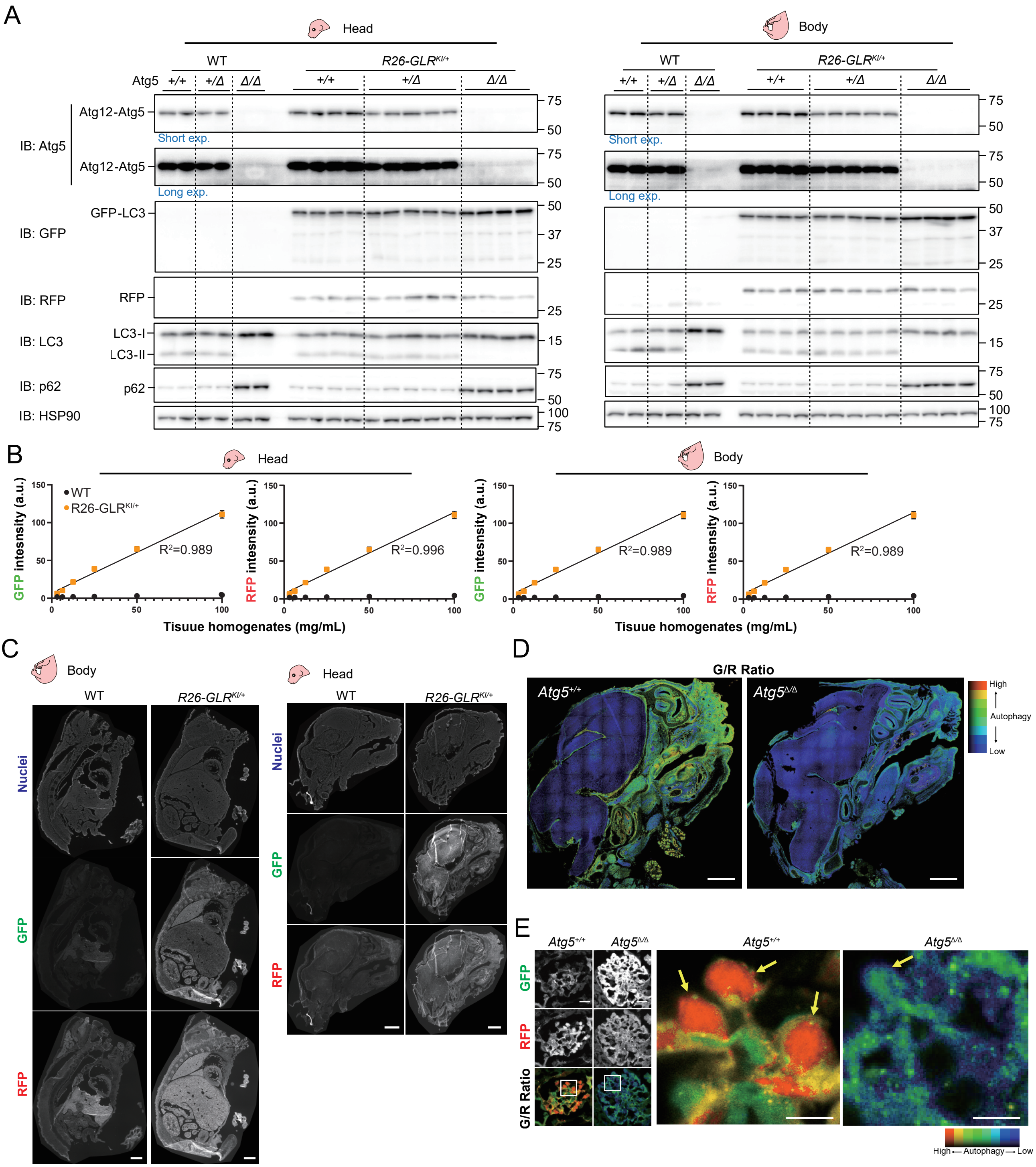

**Figure 3—figure supplement 1. Validation of R26-GLR reporter expression and Atg5 deletion in embryonic tissues.**
(A) Representative immunoblotting of head and body tissues from Atg5 wild-type (WT), heterozygous (HT), or knockout (KO) embryos with or without the GFP–LC3–RFP reporter.
(B) Soluble fractions from the head and body lysates of WT and R26-GLR embryos were serially diluted and measured using a microplate reader ($n$ = 3 per group). Data are presented as mean ± SD values. The coefficients of determination ($R^2$) of linear regressions of measurements of tissue concentration and GFP signal for R26-GLR embryos are shown.
(C) Representative tissue fluorescence images of WT and R26-GLR embryos. Scale bars, 1 mm (body) and 200 µm (head).
(D) Representative GFP/RFP ratio images of the head of *Atg5* WT and *Atg5* KO embryos. Scale bar, 200 µm.
(E) Representative GFP/RFP ratio images of kidney glomeruli of *Atg5* WT and *Atg5* KO embryos. Yellow arrows indicate podocytes. Scale bar, 100 µm.

Figure 4—figure supplement 1

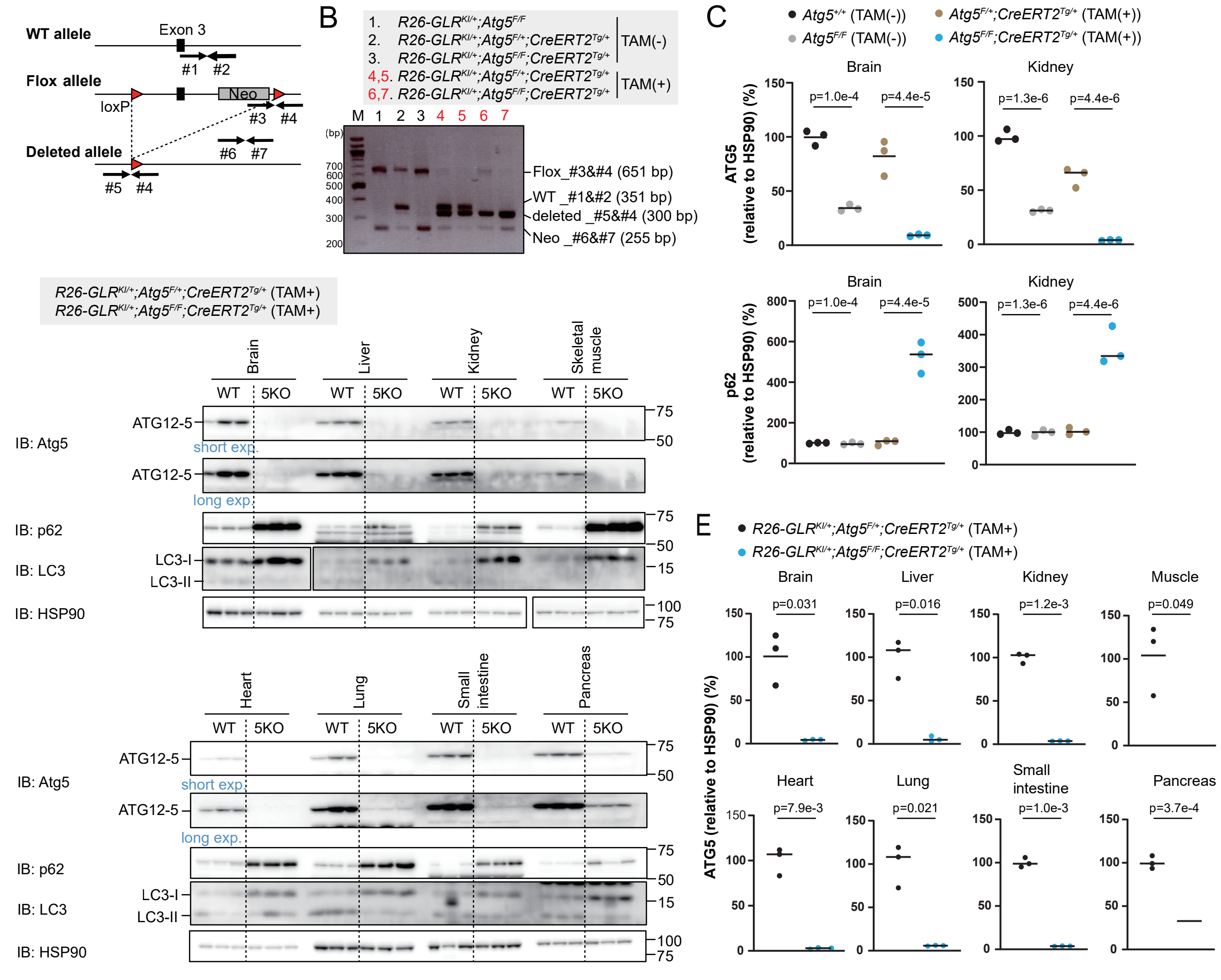


**Figure 4—figure supplement 1. Validation of tamoxifen-induced Atg5 knockout and dynamic differences in basal autophagy between embryonic and adult tissues.**

(A) Schematic illustration of Atg5 WT, flox, and deleted alleles. The positions of the PCR primers are indicated.

(B) Representative genomic PCR results of *R26-GLR*$^{KI/+}$*;Atg5*$^{flox/flox}$, *R26-GLR*$^{KI/+}$*;Atg5*$^{flox/+}$*;CreERT2*$^{Tg/+}$, and *R26-GLR*$^{KI/+}$*;Atg5*$^{flox/flox}$*;CreERT2*$^{Tg/+}$ mice with or without tamoxifen administration. Tail samples were used for PCR.

(C) Quantification of the ATG5 and p62 protein expression levels in *R26-GLR*$^{KI/+}$*;Atg5*$^{flox/flox}$, *R26-GLR*$^{KI/+}$*;Atg5*$^{flox/+}$*;CreERT2*$^{Tg/+}$, and *R26-GLR*$^{KI/+}$*;Atg5*$^{flox/flox}$*;CreERT2*$^{Tg/+}$ mice with or without tamoxifen administration. Each dot represents an individual mouse; bars indicate mean values normalized to HSP90 expression ($n$ = 3 per group). Differences between groups were analyzed by Welch's $t$-test.

(D) Representative immunoblotting of major tissues from *R26-GLR*$^{KI/+}$*;Atg5*$^{F/+}$*;CreERT2*$^{Tg/+}$ and *R26-GLR*$^{KI/+}$*;Atg5*$^{F/F}$*;CreERT2*$^{Tg/+}$ mice.

(E) Quantification of the ATG5 protein expression levels in *R26-GLR*$^{KI/+}$*;Atg5*$^{F/+}$*;CreERT2*$^{Tg/+}$ and *R26-GLR*$^{KI/+}$*;Atg5*$^{F/F}$*;CreERT2*$^{Tg/+}$ mice. Each dot represents an individual mouse; bars indicate mean values normalized to expression of HSP90; $n$ = 3 per group. Differences between groups were analyzed by Welch's $t$-test.

Figure 5—figure supplement 1

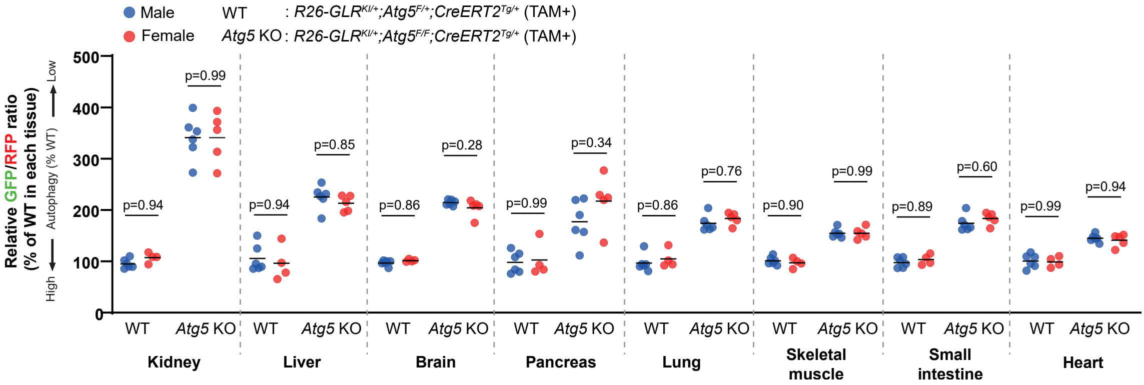


**Figure5—figure supplement 1. Evaluation of potential sex differences in basal autophagic flux across tissues.** Tissue-specific GFP/RFP ratios, as measured by microplate reader, of *R26-GLR*$^{KI/+}$*;Atg5*$^{F/F}$*;CreERT2*$^{Tg/+}$ mice, normalized to corresponding values of *R26-GLR*$^{KI/+}$*;Atg5*$^{F/+}$*;CreERT2*$^{Tg/+}$ mice (set as 100%), plotted separately for male and female mice (derived from the same dataset as in Figure 5A). Each dot represents an individual mouse; bars indicate mean values; *n* = 5–6 per group. Differences between sexes across tissues were analyzed by two-way ANOVA with Tukey's post hoc test.

Figure 6—figure supplement 1

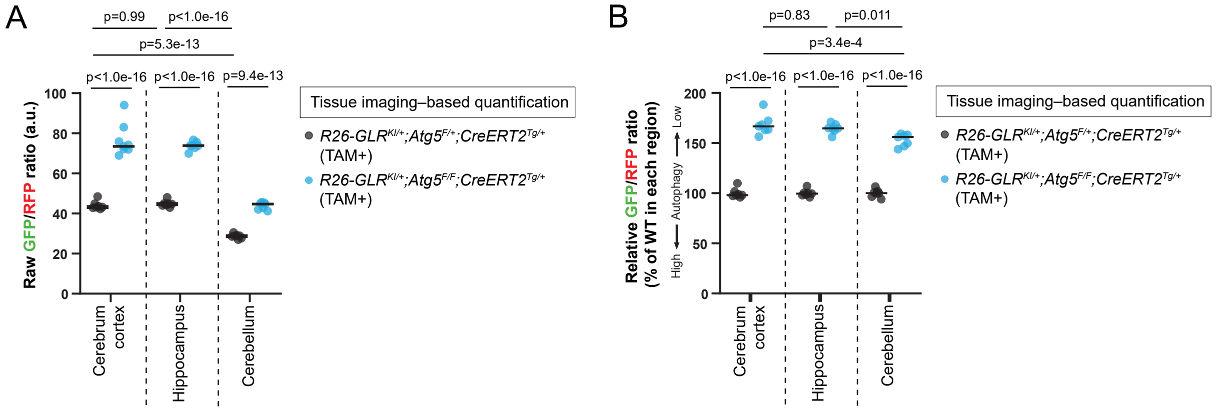


**Figure 6—figure supplement 1. Comparable levels of basal autophagic flux across brain regions.**
(A) Region-specific GFP/RFP ratios (raw values) in the brain of *R26-GLR*$^{KI/+}$*;Atg5*$^{F/+}$;*CreERT2*$^{Tg/+}$ and *R26-GLR*$^{KI/+}$*;Atg5*$^{F/F}$;*CreERT2*$^{Tg/+}$ mice measured by tissue imaging. Each dot represents an individual mouse; bars indicate mean values; *n* = 7 per group. Differences among groups were analyzed by two-way ANOVA with Tukey's post hoc test.
(B) Region-specific GFP/RFP ratios (normalized values). The data shown in (A) were normalized to values in each corresponding region of *R26-GLR*$^{KI/+}$*;Atg5*$^{F/+}$;*CreERT2*$^{Tg/+}$ mice (set as 100%). Differences among groups were analyzed by two-way ANOVA with Tukey's post hoc test.

Supplementary Table 1. List of primers used in the study

| Gene name | Forward, 5’- 3’ sequence | Reverse, 5’- 3’ sequence |
|---|---|---|
| *RFP* | CGAGGACGTCATCAAGGAGT | CTTGGCCATGTAGGTGGTCT |
| *Internal control (Atg5)* | GAATATGAAGGCACACCCCTGAAATG | GTACTGCATAATGGTTTAACTCTTGC |
| *Cre* | AGGTTCGTTCACTCATGGA | TCGACCAGTTTAGTTACCC |
| *CAG-GFP* | GCTCTAGAGCCTCTGCTAACCATGTT | TGTAGTTGCCGTCGTCCTTGAAGAAG |
| *Neo* | TGGAAGCCGGTCTTGTCGATCAGGATGATC | CAAGCTCTTCAGCAATATCACGGGTAGCC |

Supplementary Table 2. Statistical analysis of tissue-specific basal autophagic flux differences

| Sidak's multiple comparisons test | Predicted (LS) mean diff. | 95.0% CI of diff. | Summary | Adjusted P Value | Effect size (Cohen's d) |
|---|---|---|---|---|---|
| Kidney:WT vs. Kidney:5KO | -241.1 | -275.3 to -206.9 | **** | $<1.0 \times 10^{-16}$ | -11.08 |
| Kidney:5KO vs. Liver:5KO | 121.4 | 87.9 to 154.8 | **** | $<1.0 \times 10^{-16}$ | |
| Kidney:5KO vs. Brain:5KO | 131.4 | 97.9 to 164.8 | **** | $<1.0 \times 10^{-16}$ | |
| Kidney:5KO vs. Pancreas:5KO | 145.8 | 112.4 to 179.2 | **** | $<1.0 \times 10^{-16}$ | |
| Kidney:5KO vs. Lung:5KO | 162.8 | 129.4 to 196.3 | **** | $<1.0 \times 10^{-16}$ | |
| Kidney:5KO vs. Muscle:5KO | 186.4 | 152.9 to 219.8 | **** | $<1.0 \times 10^{-16}$ | |
| Kidney:5KO vs. Intestine:5KO | 191.3 | 157.8 to 224.7 | **** | $<1.0 \times 10^{-16}$ | |
| Kidney:5KO vs. Heart:5KO | 197.8 | 164.4 to 231.2 | **** | $<1.0 \times 10^{-16}$ | |
| Liver:WT vs. Liver:5KO | -119.7 | -154.0 to -85.5 | **** | $<1.0 \times 10^{-16}$ | -5.50 |
| Liver:5KO vs. Brain:5KO | 10.0 | -23.4 to 43.4 | ns | >0.99 | |
| Liver:5KO vs. Pancreas:5KO | 24.4 | -9.0 to 57.8 | ns | 0.68 | |
| Liver:5KO vs. Lung:5KO | 41.5 | 8.1 to 74.9 | ** | $1.82 \times 10^{-3}$ | |
| Liver:5KO vs. Muscle:5KO | 65.0 | 31.6 to 98.4 | **** | $8.96 \times 10^{-9}$ | |
| Liver:5KO vs. Intestine:5KO | 69.9 | 36.5 to 103.3 | **** | $4.95 \times 10^{-10}$ | |
| Liver:5KO vs. Heart:5KO | 76.4 | 43.0 to 109.8 | **** | $9.06 \times 10^{-12}$ | |
| Brain:WT vs. Brain:5KO | -109.7 | -144.0 to -75.5 | **** | $<1.0 \times 10^{-16}$ | -5.04 |
| Brain:5KO vs. Pancreas:5KO | 14.4 | -19.0 to 47.8 | ns | >0.99 | |
| Brain:5KO vs. Lung:5KO | 31.5 | -1.9 to 64.9 | ns | 0.10 | |
| Brain:5KO vs. Muscle:5KO | 55.0 | 21.6 to 88.4 | **** | $2.38 \times 10^{-6}$ | |
| Brain:5KO vs. Intestine:5KO | 59.9 | 26.5 to 93.3 | **** | $1.65 \times 10^{-7}$ | |
| Brain:5KO vs. Heart:5KO | 66.4 | 33.0 to 99.9 | **** | $3.87 \times 10^{-9}$ | |
| Pancreas:WT vs. Pancreas:5KO | -95.3 | -129.6 to -61.1 | **** | $<1.0 \times 10^{-16}$ | -4.38 |
| Pancreas:5KO vs. Lung:5KO | 17.1 | -16.4 to 50.5 | ns | >0.99 | |
| Pancreas:5KO vs. Muscle:5KO | 40.6 | 7.1 to 74.0 | ** | $2.70 \times 10^{-3}$ | |
| Pancreas:5KO vs. Intestine:5KO | 45.5 | 12.1 to 78.9 | *** | $2.90 \times 10^{-4}$ | |
| Pancreas:5KO vs. Heart:5KO | 52.0 | 18.6 to 85.4 | **** | $1.14 \times 10^{-5}$ | |
| Lung:WT vs. Lung:5KO | -78.3 | -112.5 to -44.0 | **** | $9.34 \times 10^{-12}$ | -3.60 |
| Lung:5KO vs. Muscle:5KO | 23.5 | -9.9 to 56.9 | ns | 0.77 | |
| Lung:5KO vs. Intestine:5KO | 28.4 | -5.0 to 61.8 | ns | 0.27 | |
| Lung:5KO vs. Heart:5KO | 35.0 | 1.5 to 68.4 | * | $2.79 \times 10^{-2}$ | |
| Muscle:WT vs. Muscle:5KO | -54.7 | -89.0 to -20.5 | **** | $5.48 \times 10^{-6}$ | -2.52 |
| Muscle:5KO vs. Intestine:5KO | 4.9 | -28.5 to 38.3 | ns | >0.99 | |
| Muscle:5KO vs. Heart:5KO | 11.4 | -22.0 to 44.9 | ns | >0.99 | |
| Intestine:WT vs. Intestine:5KO | -49.9 | -84.1 to -15.6 | **** | $6.24 \times 10^{-5}$ | -2.29 |
| Intestine:5KO vs. Heart:5KO | 6.5 | -26.9 to 40.0 | ns | >0.99 | |
| Heart:WT vs. Heart:5KO | -43.3 | -77.6 to -9.1 | ** | $1.28 \times 10^{-3}$ | -1.99 |

Data were analyzed using a two-way ANOVA followed by Sidak's multiple comparisons test. P values are indicated as **** $P < 0.0001$, *** $P < 0.001$,  $P < 0.01$, and * $P < 0.05$. Effect sizes (Cohen's d) are presented for WT vs. 5KO comparisons within each tissue. LS, least squares; CI, confidence interval; ns, not significant ($P > 0.05$).

| Key Resources Table | | | | |
|---|---|---|---|---|
| **Reagent type (species) or resource** | **Designation** | **Source or reference** | **Identifiers** | **Additional information** |
| strain, strain background (*Mus musculus*) | C57BL/6JJmsSlc | Japan SLC | | |
| strain, strain background (*Mus musculus*) | Atg5$^{flox/+}$ | PMID: 16625204 | RIKEN BRC: RBRC 02975 (B6.129S-Atg5$^{tm1Myok}$) | |
| strain, strain background (*Mus musculus*) | Vasa-Cre$^{Tg/+}$ | PMID: 17551945 | JAX: 018980 (B6.FVB-Tg(Ddx4-cre)1Dcas/KnwJ) | |
| strain, strain background (*Mus musculus*) | Ubc-CreERT2$^{Tg/+}$ | PMID: 18371340 | JAX: 007001 (B6.Cg- Ndor1$^{Tg(UBC-cre/ERT2)1Ejb}$/1J) | |
| strain, strain background (*Mus musculus*) | R26- STOP-GLR | This paper | RIKEN BRC: RBRC 12610 (B6J.B6N-Gt(ROSA)26Sor$^{tm1(CAG-EGFP/Map1lc3b/mRFP1)Nmz}$) | Contains STOP cassette |
| strain, strain background (*Mus musculus*) | R26-GLR | This paper | RIKEN BRC: RBRC 12611 (B6J.B6N-Gt(ROSA)26Sor$^{tm1.1(CAG-EGFP/Map1lc3b/mRFP1)Nmz}$) | STOP cassette removed |
| antibody | Rabbit polyclonal anti-LC3 | PMID: 23209294 | N/A | 1:1000 |

| | | | | |
|---|---|---|---|---|
| antibody | Rabbit polyclonal anti-LC3B | Novus Biologics | Cat#NB100-2220; RRID:AB_10003146 | 1:1000 |
| antibody | Rabbit polyclonal anti-Atg5 (C-terminal) | Sigma-Aldrich | Cat#A0731; RRID:AB_796188 | 1:1000 |
| antibody | Rabbit monoclonal anti-Phospho-p70 S6 Kinase (Thr389) (108D2) | Cell Signaling Technology | Cat#9234; RRID:AB_2269803 | 1:1000 |
| antibody | Rabbit polyclonal anti-4E-BP1 | Cell Signaling Technology | Cat#9452; RRID:AB_331692 | 1:1000 |
| antibody | Mouse monoclonal anti-HSP90 | BD Biosciences | Cat#610419; RRID:AB_397799 | 1:1000 |
| antibody | Rabbit polyclonal anti-p62 (SQSTM1) | MBL | Cat#PM045; RRID:AB_1279301 | 1:1000 |
| antibody | Rabbit polyclonal anti-Lamin A/C | Cell Signaling Technology | Cat#2032; RRID:AB_2136278 | 1:1000 |
| antibody | Rabbit polyclonal anti-GFP | Invitrogen | Cat#A-6455; RRID:AB_221570 | 1:1000 |
| antibody | Mouse monoclonal anti-RFP (clone 1G9,3G5 (mixed)) | MBL | Cat#M208-3; RRID:AB_3076166 | 1:1000 |
| antibody | Peroxidase AffiniPure F(ab')$_2$ Fragment Rabbit Anti-Mouse IgG (H+L) | Jackson Immuno Research Laboratories | Cat#315-036-003; RRID:AB_2340071 | 1:10000 |

| | | | | |
|---|---|---|---|---|
| antibody | Peroxidase-AffiniPure Goat Anti-Rabbit IgG (H+L) (min X Hu,Ms,Rat Sr Prot) | Jackson Immuno Research Laboratories | Cat#111-035-144; RRID:AB_2307391 | 1:10000 |
| antibody | Rabbit monoclonal anti-NeuN (EPR12763) | Abcam | Cat# ab177487; RRID:AB_2532109 | 1:200 |
| antibody | Goat Anti-Rabbit IgG (H+L) Highly Cross-adsorbed Antibody, Alexa Fluor 647 Conjugated | Invitrogen | Cat# A-21245; RRID:AB_2535813 | 1:500 |
| chemical compound, drug | Tamoxifen | Sigma-Aldrich | Cat#T5648 | |
| chemical compound, drug | Sunflower Seed Oil | FUJIFILM Wako | Cat#196-15265 | |
| software, algorithm | ImageJ | NIH | RRID:SCR_003070 | |
| software, algorithm | Metamorph (version 7.8.4.0) | Molecular Devices | RRID:SCR_002368 | |
| software, algorithm | GraphPad Prism (version 8.4.3) | GraphPad Software, Inc. | RRID:SCR_002798 | |
| other | | | | |
